# K48-ubiquitin-activated proteases cut-up post-ER proteins

**DOI:** 10.1101/2025.04.13.648637

**Authors:** Annabel Y Minard, Stanley Winistorfer, Liping Yu, Robert C Piper

**Author notes:** Corresponding authors (RCP).

## Abstract

Polyubiquitin chains, linked via K48 or K63 of ubiquitin, target membrane proteins in the secretory system to degradative pathways. However it’s unclear whether these linkage isomers are functionally interchangeable. Here we show that for post-endoplasmic reticulum (ER) proteins, K63-linked polyubiquitination induces multivesicular bodies (MVBs) sorting and lysosomal degradation. In contrast, K48-linked polyubiquitination induces shearing from the membrane. Substrates are cleaved by the proteasome and by two ubiquitin-activated proteases: Ddi1, a conserved cytosolic ubiquilin that generates cytosolic fragments, and Rbd2, an intramembrane rhomboid protease that produces lumenal fragments. Rbd2 localizes to Golgi/endosomes but also acts on ubiquitinated substrates at the vacuolar membrane. Ddi1’s catalytic core, the HDD-RVP domain is sufficient for ubiquitin-dependent proteolysis. It binds ubiquitin directly and its activity is amplified by auxiliary ubiquitin binding domains: an atypical UBL domain and a UBA domain. These findings demonstrate that polyubiquitin chains linked by different residues encode distinct degradative fates for post-ER proteins, and reveals two proteases that target ubiquitinated integral membrane cargo.

## Introduction

In the secretory system post-translational modification with ubiquitin (Ub) marks membrane proteins for degradation. Degrading membrane proteins is challenging since they span three regions: cytosol, lipid bilayer and lumen. This challenge is overcome by multiple mechanisms. One mechanism utilized in the endoplasmic reticulum (ER) and Golgi, involves retrotranslocating proteins, through channels formed by Ub-ligase complexes, to the cytosol for proteasomal degradation. Pathways that utilize this mechanism are known as ER-associated degradation (ERAD) and endosomal Golgi associated degradation (EGAD) ^1, 2^. A second mechanism, known as multivesicular body (MVB) sorting, degrades post-ER membrane proteins ^3^. This involves the ESCRT apparatus sorting substrates into intralumenal vesicles (ILVs), which are destroyed when MVBs fuse with lysosomes. A third mechanism may also degrade membrane proteins; in this mechanism, membrane proteins are clipped into smaller fragments for lysosomal and/or proteasomal degradation ^4^. Although clipping of membrane proteins is a well-documented cell signaling mechanism, little is known about this process in the context of Ub-dependent degradation.

How substrates are assigned to each pathway may depend not only on their localization but also on the type of ubiquitination. A feature of Ub is that it can be attached to proteins in various configurations: as a monomer or polymer, where additional Ubs are linked to one of the seven lysine residues or the M1 residue of the preceding Ub ^5^. The configurations are predicted to convey distinct signals to cellular processes and thereby function as a “ubiquitin code”. A Ub code for membrane proteins might be formed by K48- and K63-linked polyUb. These chains correlate with different degradation pathways: substrates of ERAD and proteasomal degradation are modified with K48 polyUb, whereas substrates of MVB sorting and lysosomal degradation are modified with mono-Ub and K63 polyUb ^6, 7^. Rsp5 is a major ubiquitin ligase for MVB cargo, attaching both mono-ubiquitin and K63-linked polyUb chains. Blocking its ability to form K63-linked chains results in defects in MVB sorting ^3, 7–11^. Moreover simulating its activity, by translational fusion of mono-Ub or a linear Ub chain that mimics the conformation of K63 polyUb, is sufficient for MVB sorting ^7, 12, 13^. However, whether K48 polyUb directs cargo towards the MVB pathway like K63 polyUb, or directs cargo towards a separate fate as predicted by the Ub code is unclear. This is more unclear given that both K48- and K63-polyUb can bind ESCRT proteins, which are the Ub receptors of the MVB pathway ^14, 15^.

Here we functionally test the Ub code, by leveraging targeted degradation strategies to induce K48- and K63-polyUb on post-ER membrane proteins in *S. cerevisiae*. This revealed that K63-polyUb promotes MVB sorting, whereas K48-polyUb does not.

Instead, K48-polyubiquitinated substrates undergo a distinct degradation pathway we term CUT-UP (Cleavage of Ubiquitinated Targets by Ubiquitin-activated Proteases). During this process, the K48 polyubiquitinated substrates are cut-up by three Ub-activated proteases; the proteasome and the proteases Ddi1 and Rbd2. This produces smaller fragments for proteasome and lysosome degradation. Our investigations revealed not only a Ub code for post-ER proteins, but also an alternative mechanism of degradation, and the Ub-regulated activity of two proteases.

## Methods

### Plasmids and Strains

Plasmids and yeast strains are described in Supplemental Table 1 & 2. DNA cassettes used for genome engineering yeast and constructing plasmids, were sourced from the yeast genome, plasmids or synthesized (IDT, Corallville, IA). DNA constructs were assembled using Gibson assembly cloning (NEB BioLabs, Ipswich, MA). Site directed PCR-based mutagenesis was performed using Pfu Ultra polymerase (Quick Change, Agilent Technologies, Santa Clara, CA). DNA constructs were verified by Sanger or Oxford nanopore sequencing (UIowa Genomics core (Iowa City) and Plasmidsaurus (Eugene, OR)). Yeast genomes were edited using homologous recombination. Annotated maps of key plasmids and engineered loci within the yeast strains are described (10.6084/m9.figshare.28678325.v1).

All yeast strains were confirmed for their ability to respire on ethanol/glycerol as the carbon source prior to use and storage. All *pep4* mutants were passaged until Pep4-dependent mature CPY was no longer detected as described ^16^. Strains carrying temperature-sensitive *cdc48-2* and *cdc48-3* alleles in BY4742 as described Guerriero, Weiberth and Brodsky ^17^ were confirmed to be inviable at 37°C. Strains with defined Ub^K>R^ mutations were as described in Meza Gutierrez, Simsek, Mizrak, Deutschbauer, Braberg, Johnson, Xu, Shales, Nguyen, Tamse-Kuehn, Palm, Steinmetz, Krogan and Toczyski ^18^. Briefly, in these strains all four Ub encoding genes (*UBI1*-*UBI4*) are modified to ensure a given Ub^K>R^ mutant is the sole source of Ub, with the exception of the Ub^K48R^ strain. In Ub^K48R^ strain, one of the three tandem copies of Ub in the UBI3 locus encodes wildtype Ub, providing ∼80% replacement of the Ub pool with Ub^K48R^. The proteasome mutant strains were as described in Arendt and Hochstrasser ^19^, in these strains *PRE3* and *PUP1* was deleted and in control strains wt alleles of *PRE3* and *PUP1* were reintroduced, and in catalytic inactive strains *pre3-T20A* and *pup1-T30A* were introduced. For SCF_TIR1_ assays, Skp1-TIR1 fusion protein (*S. cerevisiae* Skp1 and *O. sativa* TIR1) under the control of a *CUP1* promoter was integrated into the *PDR5* locus. Cargo targeted for SCF_TIR1_ degradation were tagged with Auxin-Inducible Degron (AID: residues 63-111 of *A. thaliana* IAA17:ADB93635). Endogenous Vps10 was modified for all degradation assays. Vps10 was tagged C-terminally with NeonGreen-AID. To N-terminally tag Vps10 with mCherry, the promotor was replaced with *MET25* promoter. For nanobody (α-GFP)-Ub-E3 ligase assays, the E3 ligase was encoded in a tetracycline/doxycline inducible low-copy plasmid. The bicistronic plasmid encoded the TetOn3G protein ^20^ constitutively expressed from a modified *TEF1*\* promoter ^21^, and terminated by both Deg1 and ADH terminator sequences to prevent read through of the second ORF. The second ORF expressed the E3 ligases from a synthetic low-background promoter derived from *PDR3* and 8 tandem tetO sites. The VHH-domain cABGFP4 nanobody was used ^22^.

### Cell Growth and Assays

Yeast growth was monitored using optical density at 600 nm (OD_600_). Cells were inoculated from a starter culture and grown overnight to an OD_600_ of 0.3-0.6 at 29°C, on an orbital shaker at 190 RPM unless otherwise stated. Synthetic complete media contained 2% glucose, yeast nitrogen base without amino acids and ammonium sulfate (Difco^TM^ Becton Dickinson, Franklin Lakes, NJ, or Y1251 Sigma Aldrich), 0.5% ammonium sulfate (Research Products International, Mount Prospect, IL), and complete supplement drop out mixture that allowed for appropriate selection (Formedium, Norfolk, UK).

For SCF_TIR1_ degradation assays, degradation was triggered by addition of 10µM CuCl_2_, to induce expression of Skp1-TIR1, and 1mM indole-3-acetic acid (IAA, Sigma-Aldrich, St. Louis, MO), to induce interaction of AID and TIR1. Cells were harvested 60 min later. Skp1-TIR1 was placed under an inducible *CUP1* promoter, since TIR1 partially induced degradation of AID tagged cargo in the absence of auxin.

For α-GFP-Ub-E3 ligase degradation assays, E3 ligase expression from a yeast TetOn3G plasmid was induced by 100 µg/ml doxycycline (Sigma-Aldrich, St. Louis, MO) and cells were harvested 60 min later.

To inhibit proteasome activity cells were treated with 100 mM MG132 (UBPbio, Dallas, TX) 30 min prior to inducing degradation. For these experiments, *PDR5* was disrupted by insertion of the *CUP1_pr_*-Skp1-TIR1 fusion gene.

Expression of mCherry-Vps10-NeonGreen-AID fusion protein was controlled by the *MET25* promoter. To avoid overexpression and mis-targeting to the vacuole, 10 mg/ml methionine (Alfa Aesar, Ward Hill, MA) was included in the growth medium.

Expression of mCherry-Vps10-sfGFP-DHRF-AID was controlled by a modified TEF1 promoter (*TEF1*\*_pr_) and tetracycline-inducible riboswitch ^23, 24^. Expression was blocked with the addition of 0.5mM tetracycline (Research Products International, Mount Prospect, IL).

### Immunoblotting

Whole-cell extracts were prepared by resuspending pelleted (11,000 xg for 30 s) cells in 0.2 M NaOH, repelleting, and resuspending in SDS-lysis buffer (8 M urea, 5% SDS, 10% glycerol, 50 mM Tris-HCl pH 6.8, 2.5% 2-mercaptoethanol, 0.02% bromophenol blue) at a concentration of 10 OD_600_ units/ml. The samples were heated at 95°C for 1 min immediately before being loaded onto 4–12% gradient gels (ExpressPlus™, Genscript, Piscataway, NJ) for SDS-PAGE, followed by transfer to 0.22 µm nitrocellulose membranes. Primary antibodies and HRP-conjugated secondary antibodies are listed in Supplemental Table 3. Chemiluminescent signals were generated using SuperSignal^TM^ West Femto or Pico PLUS Substrate (Thermo Fisher, Waltham, MA), that were recorded with a FluorChem^TM^ 8800 (Alpha Innotech, San Leandro, CA) or an iBright^TM^ Imaging System (Thermo Fisher, Waltham, MA). Densitometry analysis was performed in Fiji ^25^.

### Microscopy

Yeast were concentrated by centrifuging at 2,000 xg for 2 min, transferred to microscope slides, and imaged at room temperature. Images were acquired using an Olympus fluorescence BX60 microscope equipped with UPlanSApo100×/1.40 oil objective and Hamamatsu Orca-R2 digital camera (Hamamatsu, JP) controlled by iVision software (BioVision Technologies).

### Subcellular Fractionation

Cell fractions were collected from pelleted yeast that were washed twice in ice cold PBS containing protease inhibitors (cOmplete EDTA-free and Pefabloc (Roche, Basel, CH) and lysed using a OneShot cell disruptor (Constant Systems, Daventry, UK) at 35 kPsi. Lysate was centrifuged at 5,000 xg for 10 min at 4°C, to generate a post nuclear supernatant, that was subsequently centrifuged at 200,0000 xg for 1 h, generating a soluble cytosolic fraction and crude membrane pellet.

### NMR

Recombinant proteins were expressed in *E. coli* BL21(DE3) strain upon induction with 0.5 mM IPTG at 18°C for 20 h in LB media supplemented with antibiotic and 0.1% glucose. Cells were lysed in ice cold PBS containing protease inhibitors (cOmplete EDTA-free and Pefabloc (Roche, Basel, CH) and lysed using a OneShot cell disruptor (Constant Systems, Daventry, UK) at 25 kPsi. 6xHis tagged proteins were purified using Talon Co^2+^ affinity resin (Takara Bio, USA) and eluted with 150 mM Imidazole in PBS buffer pH 7.8, and equilibrated to 50 mM NaCl, 40 mM NaPO4, pH 6.95. HSQC ^15^N/^1^H spectra of ^15^N-Ub (30µM) in presence and absence of recombinant Ddi1 protein domains were collected at 25 °C on a Bruker Avance II 800 MHz spectrometer and analyzed with SPARKY (T. D. Goddard and D. G. Kneller, SPARKY 3, UCSF, CA) and NMRView (One Moon Scientific, Westfield, NJ). Chemical shift perturbations were measured by comparing peak positions to ^15^N-Ub alone using ((0.2ΔN^2^ + ΔH^2^)^1/2^) to map residues in and near binding interfaces on Ub. The bound fraction in titration experiments was calculated by measuring the difference in the peak intensity in the absence (free form) and presence (bound form) of the Ub-binding protein, then divided by the peak intensity of the free form. These data were then fitted to a standard quadratic equation using GraphPad Prism (GraphPad Software). The standard deviation from data fitting is reported.

## Results

### K48 and K63 Ub-ligases target a post-ER protein for degradation via different pathways

To investigate the effect of K48- or K63-linked polyubiquitination on post-ER proteins, we constructed a panel of synthetic Ub-ligases that preferentially form either polyubiquitin chain and targeted them to a post-ER protein. The CPY receptor, Vps10, was selected as a model substrate because it has simple Type-1 topology, is stable over a 2h period, and traffics between TGN and late endosomal compartments, where we have previously shown it is susceptible to MVB sorting when fused to mono-Ub ^13, 26-^ 28.

The panel of Ub-ligases consisted of a HECT, RING and SCF-type Ub-ligases (Figure 1A-B). The HECT and RING type Ub-ligases were constructed by fusing an α-GFP nanobody to the catalytic domain of an E3 Ub-ligase that preferentially forms either K48 or K63 polyUb (Figure 1A) ^29^. Expression of the α-GFP Ub ligases was controlled by a modified TetOn 3G system, which produced the fusion proteins within 30 min of doxycycline addition. Endogenous Vps10 was tagged with GFP to recruit the ligases. To stimulate K63-ubiquitination we used the HECT domain of Rsp5 ^9^ and the RING domain of Pib1 ^30, 31^. To stimulate K48-ubiquitination we a used a RSP5-E6AP chimeric HECT domain and a dimer of the RNF4 RING domain ^32^. In the chimera, the C-lobe of Rsp5 HECT domain was swapped for the C-lobe of E6AP HECT domain, this configuration was previously shown to switch Rsp5 activity from producing K63 to K48 polyUb ^33^.

**Figure 1.**
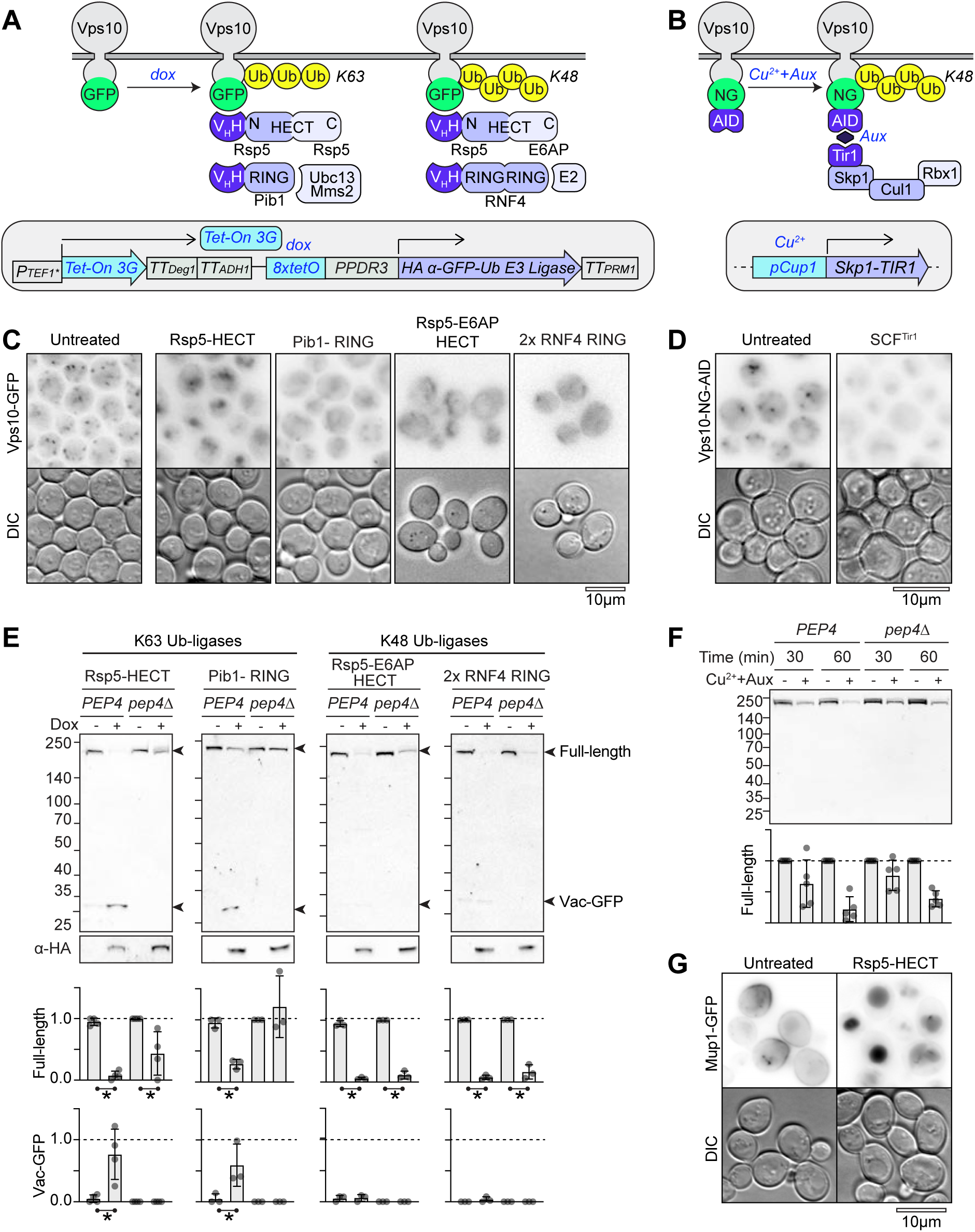
Ubiquitin-linkage specifies distinct degradation pathways for post-ER membrane protein. **A-B)** Schematic of inducible ubiquitination systems. **A)** K63-Ub-ligases were engineered by fusing a GFP nanobody to the RING domain of Pib1 or HECT domain of Rsp5, K48-Ub ligases were engineered by fusing a GFP nanobody to a chimera of the Rsp5 and E6AP HECT domains or to two RNF4 RING domains. GFP nanobody Ub-ligases were placed under the TET-On promoter and induced with doxycycline to trigger ubiquitination of endogenous Vps10 that was tagged with GFP. **B)** K48-polyubiquitination was also stimulated using the TIR1-AID system. Vps10 was tagged with NeonGreen (NG) and AID. TIR1 was integrated in the *PDR5* locus and placed under the *CUP1* inducible promoter. **C)** Fluorescence localization of Vps10-GFP after 1h expression of nanobody ligase indicated; DIC=Differential Inference Contrast. **D)** Localization of Vps10-NG-AID 1h after induction of TIR1-dependent ubiquitination with copper and auxin. **E)** Immunoblot of Vps10-GFP after 1h of doxycycline treatment to induce GFP nanobody in *PEP4* and *pep4*Δ cells. Levels of full-length Vps10-GFP and vacuolar-processed GFP were quantified (n ≥3, *p<0.05). **F)** Immunoblot of Vps10-NG-AID after 1h induction of TIR1 with copper and auxin, in *PEP4* and *pep4*Δ cells. Levels of Vps10-NG-AID were quantified (n ≥3, *p<0.05). **G)** Micrograph of Mup1-GFP upon induction of Rsp5-HECT Ub ligase for 1h. Bar=10μm.

To complement these ligases we also used the TIR1-AID system. TIR1, an F box protein from plants, combines with endogenous proteins to form a Skp1-Cullen-Fbox (SCF^TIR1^) K48-polyUb ligase complex (Figure 1B) ^34–37^. TIR1 interacts with the auxin-inducible degron in the presence of the auxin molecule, indole-3-acetic acid (IAA).

However, because we observed low activity in the absence of IAA, we placed TIR1 under a copper inducible promoter. To implement this system, endogenous Vps10 was tagged with neon green (NG) and a minimal auxin-inducible degron (AID) ^38^, and TIR1 was fused to Skp1 and integrated into the *PDR5* locus under a copper inducible *CUP1* promoter.

By microscopy (Figure 1C-D) and immunoblot analysis (Figure 1E-F), we observed that the Ub-ligases induced Vps10 degradation using one of two different mechanisms. The Rsp5-HECT and Pib1-RING degraded Vps10-GFP while producing Vac-GFP, a remanent of GFP produced by vacuolar proteases ^39^. In *pep4*Δ mutants which lack active vacuolar proteases, Vps10-GFP was largely stabilized and no Vac-GFP was produced. These data indicated that K63-polyUb is sufficient to engage MVB sorting.

Similarly, we found that the Rsp5-HECT could also sort a plasma membrane localized transporter, Mup1-GFP, into the vacuole (Figure 1G). In contrast the K48-Ub ligases Rsp5-E6AP HECT and RNF4-RING induced Vps10-GFP degradation in *PEP4* and *pep4*Δ cells, without increasing Vac-GFP (Figure 1E). Similarly, SCF^TIR1^ induced Vps10-NG-AID degradation in *PEP4* and *pep4*Δ cells, without increasing Vac-NG (Figure 1F).

By microscopy (Figure 1C-D) and immunoblot analysis (Figure 1E-F), we observed that the Ub-ligases induced Vps10 degradation using one of two different mechanisms. The Rsp5-HECT and Pib1-RING degraded Vps10-GFP via the MVB pathway. They produced Vac-GFP, a remanent of GFP left by vacuolar proteases ^39^, and in *pep4*Δ mutants which lack active vacuolar proteases, Vps10-GFP was largely stabilized and no Vac-GFP was produced. We verified that K63-polyUb is sufficient to engage MVB sorting, by observing that Rsp5-HECT could also sort a plasma membrane localized transporter, Mup1-GFP, into the vacuole (Figure 1G). In contrast the K48-Ub ligases Rsp5-E6AP HECT and RNF4-RING induced Vps10-GFP degradation in *PEP4* and *pep4*Δ cells, without increasing Vac-GFP (Figure 1E). Similarly, SCF^TIR1^ induced Vps10-NG-AID degradation in *PEP4* and *pep4*Δ cells, without increasing Vac-NG (Figure 1F).

These data show that K48- and K63-Ub ligases engaged distinct degradative pathways, thereby demonstrating a functional Ub code for post-ER proteins and the selectivity of the ESCRT/MVB pathway for K63-ubiquitinated cargo.

### K48-ubiquitination of post-ER Vps10 induces shearing

We next investigated how the post-ER reporter protein, Vps10, was degraded by K48-Ub ligases. For these studies we focused on the degradation of Vps10-NG-AID by the TIR1-AID system. Having ruled out degradation *via* the MVB pathway, we examined the role of proteasomes. To inhibit the proteasome we targeted all three of its proteolytic sites. The trypsin-like activity of the Pup1/β2 subunit and caspase-like activity of Pre3/β1 subunit were inactivated by using a *pup1-T30A pre3-T20A* double mutant ^19, 40^. The wildtype alleles, *PUP1 PRE3*, were reintroduced in a control strain for this mutant. The chymotrypsin-like activity of the Pre2/β5 subunit was inhibited by deleting Pdr5, a drug efflux pump, and treating cells with MG132 ^41^. All of these conditions were necessary to block cell growth or degradation of a cytosolic reporter protein (NeonGreen fused to AID) (Supplemental Figure 1A). In addition, *PEP4* was deleted to inactivate vacuolar proteases.

Immunoblots of NG revealed that inhibiting the proteasome and vacuolar proteases did not prevent degradation of Vps10-NG-AID in response to SCF^TIR1^ ubiquitination (Figure 2A). However, C-terminal fragments of Vps10-NG-AID accumulated upon proteasome inhibition. The sizes of the C-terminal fragments, which ranged between 40 kDa and 70 kDa by SDS-PAGE, were consistent with cleavage of Vps10-NG-AID in the juxtamembrane region of the cytosolic domain, which is predicted to be 52 kDa. This indicated that the cytosolic portion of Vps10-NG-AID was cleaved by a protease, which is activated by Ub, and that the cytosolic portion was degraded by the proteasome.

**Figure 2.**
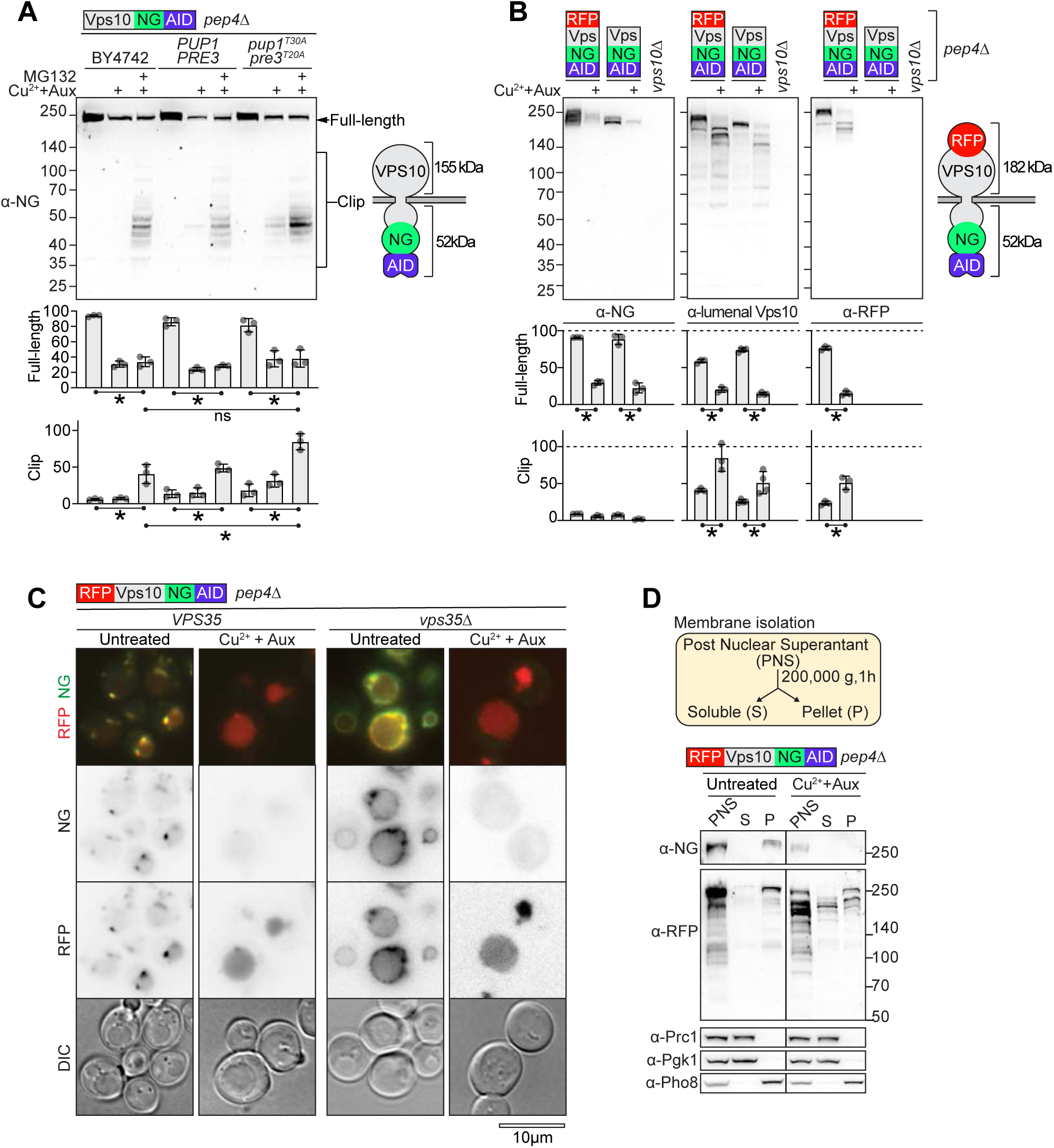
Vps10 is sheared after K48-ubiquitination. **A)** Immunoblot analysis of Vps10-NG-AID degradation in BY4742, or cells harboring wildtype *PUP1 PRE3* or catalytically inactive *pup1-T30A pre3-T20A* alleles. Cells were treated with MG132 for 30 min, followed by 1 h of copper and auxin to induce SCF^Tir1^ mediated ubiquitination. The levels of full-length Vps10-NG-AID and clipped degradation products were quantified and graphed (n ≥3, *p<0.05). **B)** Immunoblot analysis of the degradation of the lumenal or cytosolic domain of Vps10-NG-AID or RFP-Vps10-NG-AID after SCF^Tir1^-ubiquitination using antibodies detecting NG, RFP, and Vps10 (lumenal residues 307-696). Levels of full-length and clipped products were quantified and graphed (n ≥3, *p<0.05). **C)** Localization of RFP-Vps10-NG-AID in wildtype (*VPS35*) and *vps35*Δ cells before and after ubiquitination by SCF^Tir1^. **D)** Post nuclear supernatant (PNS), soluble (S), and membrane-associated (P) fractions blotted for RFP-Vps10-NG-AID as shown. **(A-E)** All cells were *pep4*Δ*pdr5*Δ. Bar=10μM.

We next tracked the fate of the lumenal domain of Vps10-NG-AID, which spans residues 1-1372, using an antibody raised against residues 307-696 of Vps10 ^27^. A truncated N-terminal fragment of Vps10-NG-AID accumulated in a *pep4*Δ strain, but not in a *PEP4* strain (Supplemental Figure 1B). The size of the fragment, which was between 140 kDa marker and full-length Vps10-NG-AID (approximately 250 kDa) by SDS-PAGE, was also consistent with cleavage of Vps10-NG-AID in the juxtamembrane region. These data indicated that the lumenal portion of Vps10-NG-AID, was delivered to and degraded in the vacuole.

Because the α-Vps10 antibody detected non-specific bands, we inserted mCherry (RFP) between the signal sequence and lumenal domain of Vps10-NG-AID. α-RFP antibodies also detected a truncated fragment of RFP-Vps10-NG-AID after SCF^TIR1^ activation, the product ran between 140 kDa and full-length RFP-Vps10-NG-AID (Figure 2B). Fluorescence microscopy confirmed that the RFP-tagged lumenal fragment redistributed from endosomes to the vacuole lumen, while the NG-tagged cytosolic domain disappeared (Figure 2C).

The localization of the RFP-tagged lumenal domain to the vacuole is expected after degradation of the cytosolic domain, since the cytosolic domain contains retrieval signals recognized by the Retromer complex. However, loss of these retrieval signals should cause Vps10 to localize to the limiting vacuolar membrane, rather than the lumen in a *pep4*Δ mutant ^26, 27, 42^. We confirmed this by examining the localization of RFP-Vps10-NG-AID in a *vps35*Δ mutant, which lacks Retromer activity (Figure 2C, Supplemental Figure 1C). In *vps35*Δ mutants, RFP-Vps10-NG-AID indeed localized to the vacuolar limiting membrane, but following SCF^TIR1^-mediated ubiquitination, RFP was released into the lumen, while NG disappeared. This indicated that the Ub-activated protease severed the lumenal domain from the membrane and that it functions in both TGN/endosomal compartments and the vacuolar membrane. We confirmed that the lumenal domain became soluble upon SCF^TIR1^-mediated ubiquitination, by isolating soluble and membrane fractions from cells expressing RFP-Vps10-NG-AID using high-pressure lysis, high-salt extraction and ultracentrifugation (Figure 2D). Full-length RFP-Vps10-NG-AID was detected in the membrane fraction, and in the soluble fraction a smaller RFP-tagged fragment was detected that increased in abundance after ubiquitination. Thus, these results demonstrate that Vps10 targeted by SCF^TIR1^ is cleaved by a Ub-activated proteases. Its cytosolic domain is degraded by the proteasome, and the lumenal domain is severed from the membrane and degraded in the vacuole.

### Ddi1 is a Ub-dependent protease that cleaves the cytosolic domain of K48-polyubiquitinated Vps10

Next, we screened for mediators of Vps10-NG-AID shearing. We searched the MEROPS database for proteases, or orthologs of proteases, that were reported to have access to endosomes, be involved in protein quality control, and that were not proteasome subunits or deubiquitinases. Of the 122 known and putative proteases, we selected 10 that met these criteria: Yps7, Ape4, Ydr415c, Ste24, Mca1, Rim13, Yps5, Ypf1, Rbd2, and Ddi1 ^43–46^. In addition to proteases, we screened the Dsc retrotranslocation complex, which operates the EGAD pathway. Although EGAD is not known to involve protein shearing, it is the only known pathway for proteasomal degradation of endosomal and Golgi substrates in *S. cerevisiae* ^2^. The Dsc complex subunits we tested were Tul1, Vld1, Gld1, Dsc3, Ubx3, Dsc2 ^47^ and Cdc48. Strains in which these genes were deleted were obtained from the yeast deletion collection, except for a *dsc2*Δ mutant, which we generated independently, and strains *carrying cdc48-2* or *cdc48-3*, which are temperature-sensitive alleles of the essential *CDC48* gene. Each strain was modified by integrating the *CUP1*-Skp1-Tir1 cassette into the *PDR5* locus, and tagging endogenous Vps10 with NG-AID. Vps10-NG-AID degradation with and without MG132 was assessed by immunoblot analysis (Supplemental Figure 2).

We found that only a *ddi1*Δ mutant exhibited a defect in Vps10-NG-AID degradation. Loss of full-length Vps10-NG-AID was still evident in *ddi1*Δ cells, however far less cleaved cytosolic fragments were produced under proteasome inhibition (Figure 3A). This was confirmed in the *pup1-T30A, pre3-T20A pep4*Δ strain, which was inactivated of lysosomal and proteasomal proteases. In this strain, high levels of cleaved cytosolic fragments were observed upon proteasome inhibition with MG132, yet in a *ddi1*Δ mutant, almost all the fragments were eliminated. Loss of full-length Vps10-NG-AID was still observed in this *ddi1*Δ mutant strain, indicating Ddi1 is responsible for forming the clipped cytosolic fragment of Vps10-NG-AID, but additional mechanisms degrade Vps10-NG-AID.

**Figure 3.**
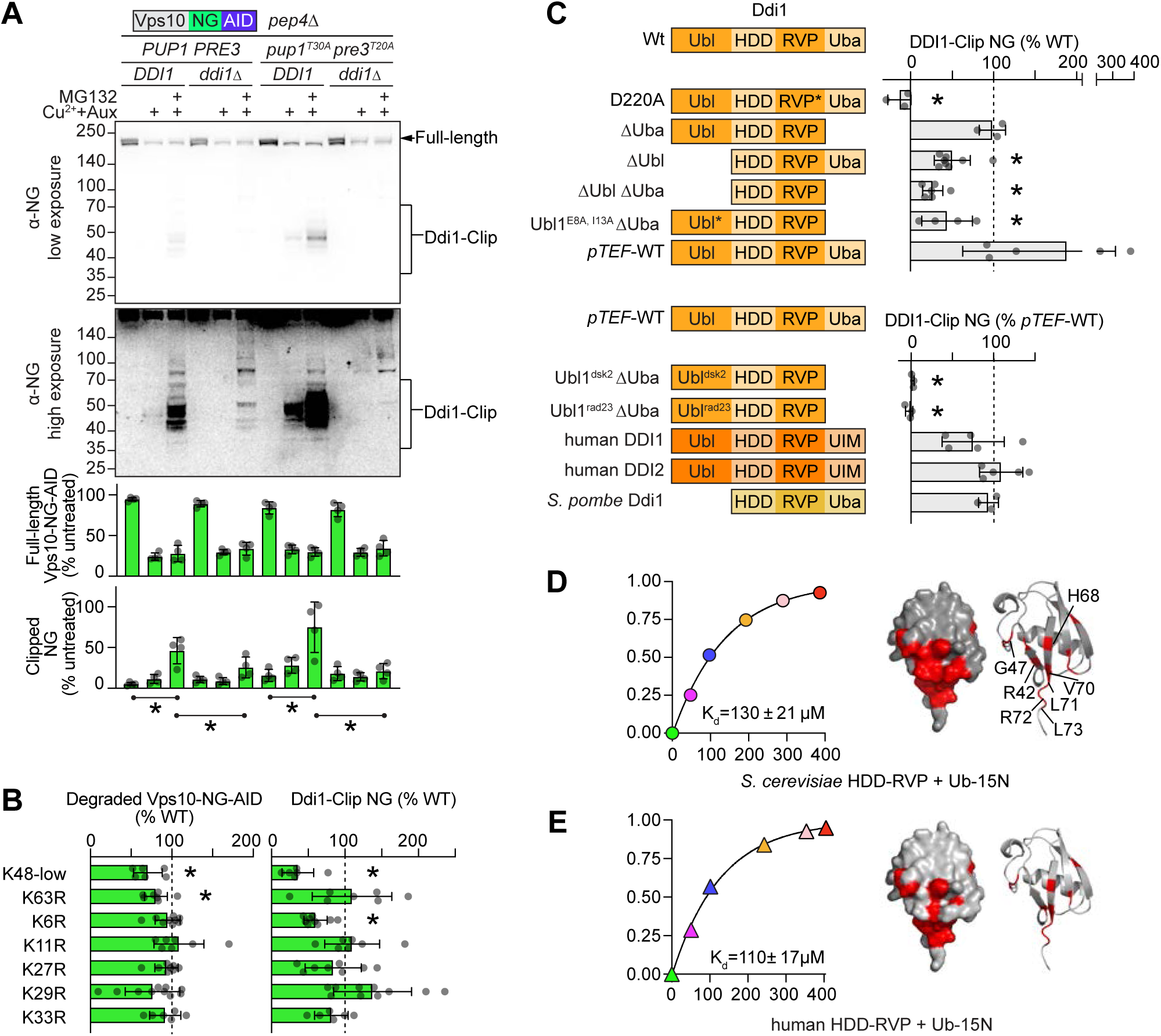
K48-ubiquitinated Vps10 is cleaved by Ddi1 **A)** Immunoblot analysis of degradation of Vps10-NG-AID after SCF^Tir1^-ubiquitination, in cells with (*DDI1I*) or without (*ddi1*Δ) Ddi1, and with wildtype (*PUP1 PRE3*) or catalytically inactive (*pup1-T30A pre3-T20A*) proteasome alleles. Full-length and clipped forms of Vps10-NG-AID were quantified and graphed (n ≥3, *p<0.05). **B)** Quantification of full-length and clipped forms of Vps10-NG-AID produced in strains carrying the indicated K>R Ub as their sole source of Ub, except for K48R Ub, which also carries low levels of wildtype Ub to sustain viability (n ≥3, *p<0.05). **C)** Quantification of clipped Vps10-NG-AID after MG132 treatment and SCF^Tir1^ induction in *pup1-T30A pre3-T20A ddi1*Δ cells expressing HA-epitope tagged Ddi1 variants (n ≥3, *p<0.05). *Upper panel,* Ddi1 constructs lacking Ubl, Uba or both domains were expressed from the *DDI1* promoter. *Lower panel,* Ddi1 constructs with Ubl domain replaced by Ubl of Dsk2 or Rad23, and human DDI1, DDI2, and S. pombe Ddi1 were expressed from a *TEF1* promoter. **D-E)** HSQC analysis of 30µM ^15^N-Ub in the presence of increasing concentrations of recombinant HDD-RVP domains from *Sc*Ddi1 **(D)** and human DDI2 **(E)**. **D-E)** *Left,* Fraction of ^15^N-Ub bound as a function of HDD-RVP concentration (µM). *Right,* Residues of Ub that underwent the largest chemical shift perturbation are mapped in red onto the structure of Ub (PDB: 1UBQ).

Ddi1 is one of three highly conserved ubiquilins, along with Rad23 and Dsk2. Ubiquilins are cytosolic proteins that contain a Ub-associated domain (UBA) domain, which binds ubiquitinated substrates ^48^, and a Ub-like (UBL) domain, that binds Ub receptors on the proteasome ^49^. Ubiquilins are thought to shuttle ubiquitinated substrates to the proteasome. Ddi1 is distinguished by a retroviral aspartyl protease-like (RVP) domain and a helical domain, dubbed the helical domain of Ddi1 (HDD). Little is known about the proteolytic substrates of Ddi1, however *in vitro* Ddi1 was shown to cleave substrates linked to K48-polyUb ^50^. Human Ddi2 also acts as a Ub-activated protease, with two known substrates; the integral membrane proteins Nrf2 and AMOT are cleaved by Ddi2 after they are ubiquitinated to form transcription factors from their cytosolic tails ^51, 52^.

We found that HA epitope-tagged Ddi1 rescued cleavage of Vps10-NG-AID in *ddi1*Δ strains, however, a catalytically dead Ddi1 (D220A) did not (Figure 3C), demonstrating that Ddi1 proteolytic activity mediated cleavage of Vps10-NG-AID.

Next we tested the requirement for K48-polyUb and other Ub linkages in Ddi1-dependent cleavage, by using a panel of mutants expressing a lysine to arginine mutation at each of Ub residues ^18^. Because K48 Ub is essential ^53^, the K48R Ub strain included one copy of wildtype K48 Ub resulting in ∼20% wildtype Ub expression ^18^. In this K48R strain, degradation of Vps10-NG-AID and formation of the Ddi1-dependent cytosolic fragment were decreased, indicating that these processes are K48 polyUb-dependent (Figure 3B). Strains carrying K6R, K11R, K27R, K29R K33R, or K63R as their sole form of Ub, all degraded Vps10-NG-AID and formed Ddi1-dependent cytosolic fragments, which demonstrated that these polyUb-linkages are not required. However, Vps10-NG-AID degradation was slightly impaired in the K63R Ub mutant, as was formation of the Ddi1-dependent cytosolic fragment in the K6R Ub mutant, potentially indicating a minor direct or indirect role for these other linkages.

Ddi1 contains two Ub binding domains that might allow it to recognize ubiquitinated Vps10-NG-AID as a substrate: a UBA domain and an atypical UBL domain which, unlike most UBL domains, can bind Ub ^45, 54^. We introduced GFP-tagged Ddi1 variants lacking either the UBA domain, the UBL domain, or both (HDD-RVP) into a *ddi1*Δ mutant and assessed cleavage of the cytosolic domain of Vps10-NG-AID (Figure 3C). We found that deletion of the UBL but not the UBA domain impaired Ddi1 activity, however Ddi1 activity was most compromised in the construct deleted of both the UBL and UBA domain. This indicated that the UBA domain is involved in Ub-dependent processing but is redundant in the presence of the UBL. Interestingly, the construct lacking both UBL and UBA domains (HDD-RVP), retained some Ub-dependent activity compared to the catalytic inactive Ddi1 (D220A). These data were consistent with the prior *in vitro* study of Ddi1 ^50^ that defined the HDD-RVP domain as the functional core of the Ub-dependent protease.

We investigated whether Ddi1-dependent proteolysis required the UBL’s ability to bind Ub or the proteasome. To investigate this, we replaced Ddi1’s UBL domain with UBL domains not able to bind Ub. In one construct, E8 and I13 in the Ddi1 UBL domain were mutated to alanine to prevent Ub-binding ^54^. In two other constructs the Ddi1 UBL domain was replaced with those of Dsk3 or Rad23, which do not bind Ub but would still mediate proteasome binding. Ddi1 activity was compromised by all the UBL replacements (Figure 3C). Therefore, the UBL domain contributes to Ddi1-activity by binding Ub rather than the proteasome.

The Ub-binding domains varies across species: human Ddi1 and Ddi2 have N-terminal UBL domains and C-terminal UIM (Ub-interaction motif) domains that bind Ub ^55^, whereas *S. pombe* Ddi1 (Mud1), has a C-terminal UBA domain that binds K48 polyUb ^56^. We found that these orthologs complemented formation of Ddi1-dependent cytosolic fragments in *ddi1*Δ mutants (Figure 3C), although clip pattern was slightly different, suggesting different architectures of Ddi1 may affect substrate positioning in the proteolytic site (Supplemental Figure 3).

Since the HDD-RVP construct, lacking both UBL and UBA domains, retained Ub-dependent proteolytic activity, we predicted that it can recognize Ub directly. Indeed, NMR HSQC experiments with ^15^N-Ub revealed that the HDD-RVP domain of Sc.Ddi1 and human Ddi2 bound mono-Ub with a Kd of ∼100µM, engaging the hydrophobic patch on Ub used by a wide variety of Ub-binding modules ^57^ (Figure 3D-E, Supplemental Figure 4). The HDD domain was required for Ub binding, since no Ub-binding could be detected by the RVP alone, either from *S. cerevisiae* Ddi1 or human Ddi2. In addition, we found that mutating conserved residues I183 or D184 in *S. cerevisiae* Ddi1 to alanine prevented proteolysis to a similar extent as catalytic inactive mutant (D220A) underscoring a critical role of the HDD domain (Supplemental Figure 3).

Collectively, these data demonstrate that Ddi1 is a K48-Ub dependent protease that can operate outside of the proteasome to target membrane proteins. It utilizes a central HDD-RVP catalytic core as its Ub-activated catalytic center, which can be enhanced by additional auxiliary Ub-binding domains.

### Rbd2 is a Ub-dependent protease that cleaves the lumenal domain of K48-polyubiquitinated Vps10

Our data showed that ubiquitinated Vps10 was targeted by another Ub-dependent protease, also responsible for generating a soluble N-terminal lumenal fragment (Figure 2D). Following the lumenal domain of RFP-Vps10-NG-AID by immunoblotting for RFP, revealed that upon SCF^TIR1^ ubiquitination two RFP-tagged products were formed that ran between full-length RFP-Vps10-NG-AID and the 140 kDa marker (Figure 4A). The larger but not smaller band was sensitive to proteasome inhibition: its formation was inhibited by MG132 in a BY4742 strain or in the proteasome control strain (*PUP1 PRE3*); and was eliminated, even without MG132, in a *pup1-T30A, pre3-T20A* mutant. In contrast, the smaller band formed in a *pup1-T30A, pre3-T20A ddi1*Δ strain, and thus was produced by an unknown protease.

**Figure 4.**
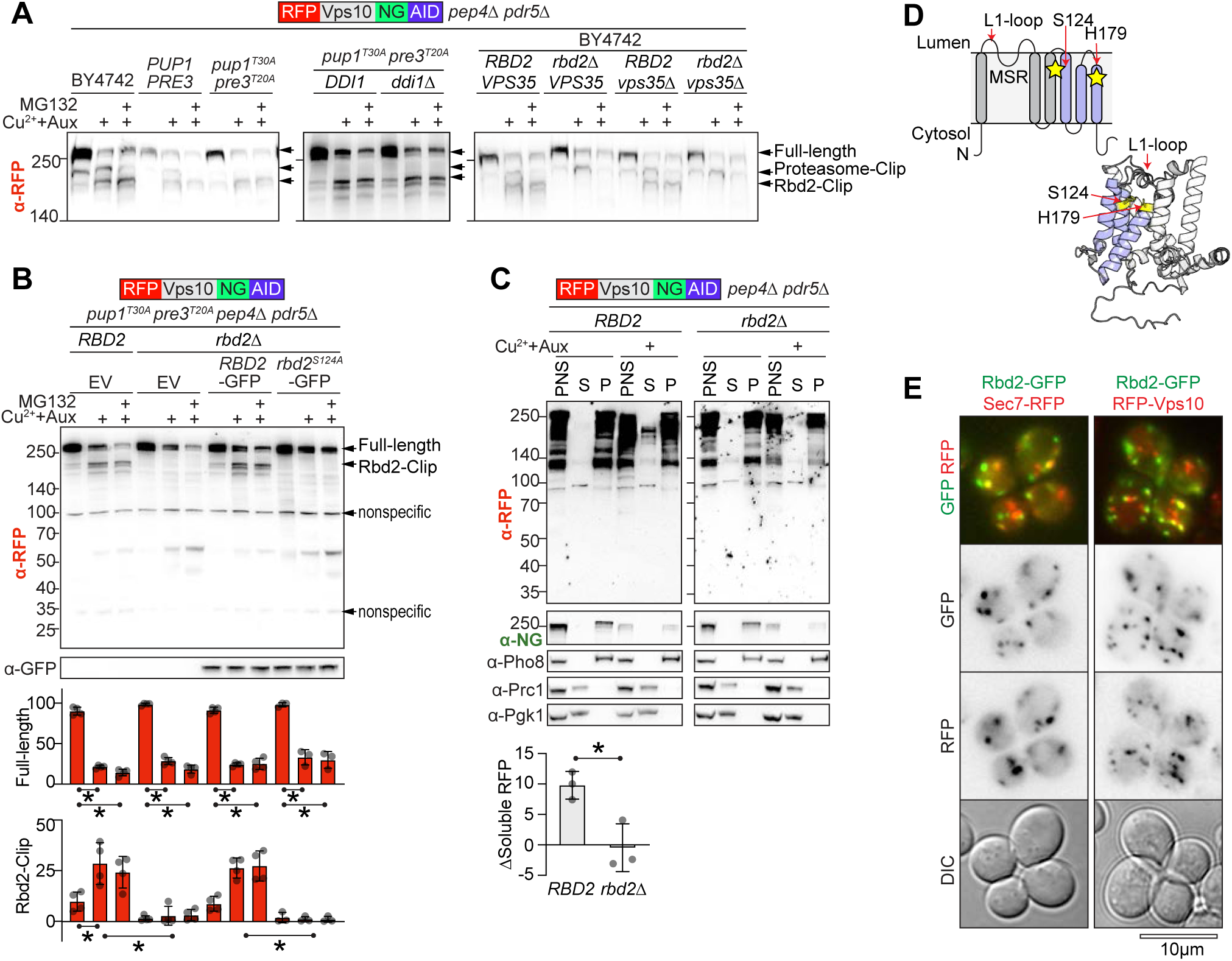
Rbd2 is a Ub-dependent protease. **A)** Proteasome and Rbd2-dependent degradation products detected by α-RFP immunoblot analysis of the lumenal domain of RFP-Vps10-NG-AID after SCF^Tir1^-directed ubiquitination. **B)** Immunoblot analysis of Rbd2-dependent degradation products in *pup1-T30A, pre3-T20A rbd2*Δ cells expressing catalytic active and inactive (S124A) Rbd2-GFP. The level of full-length and Rbd2-dependent degradation products were quantified and graphed (n ≥3, *p<0.05). **C)** Immunoblot analysis of post nuclear supernatant (PNS), soluble (S), and membrane-associated pellet (P) fractionated from BY4742 *RBD2* and *rbd2*Δ cells before and after inducing SCF^TIR1^-directed ubiquitination RFP-Vps10-NG-AID. The change in amount of soluble RFP-tagged fragments was quantified (n ≥3, *p<0.05). **D)** *Top,* schematic of the six transmembrane domains that comprise the rhomboid fold of Rbd2; *bottom,* predicted Alpha fold structure (A0A816BH90). Catalytic sites are highlighted in yellow. **E)** Micrographs of Rbd2-GFP co-expressed with Sec7-RFP or RFP-Vps10. Bar=10μM.

Rbd2 stood out as a likely candidate since it is a rhomboid protease localized to Golgi and endosomes ^58^. Although *S. cerevisiae* Rbd2 has no known substrates, evidence from its homologs—*S. pombe* Rbd2 and the ER-localized mammalian rhomboid protease RHBDL4—suggested its potential as a Ub-dependent protease ^46, 59, 60^. These homologs recognize ubiquitinated substrates *via* domains that are not conserved in *S. cerevisiae*: RHBDL4 contains a Ub-interaction-motif, and both RHBLD4 and *S. pombe* Rbd2 have SHP box domains, which recruit the Ub-adaptor protein Cdc48/p97 ^46, 60^.

Loss of Rbd2 in BY4742 cells eliminated the smaller RFP-tagged degradation fragment (Figure 4A). In a *pup1-T30A, pre3-T20A* mutant lacking Rbd2, neither the small or large RFP-tagged degradation fragments formed (Figure 4B). In BY4742 *vps35*Δ cells, where RFP-Vps10-NG-AID is relocalized to the limiting membrane of the vacuole, we observed RFP-tagged fragments that were proteasome-dependent and Rbd2-dependent (Figure 4A). Fractionation experiments demonstrated that the soluble RFP-tagged degradation fragment was a product of Rbd2, since it was not formed in *rbd2*Δ mutants (Figure 4C).

Rbd2 is a predicted rhomboid protease with six transmembrane domains, sharing a topology with the well-characterized bacterial GlpG rhomboid protease ^61, 62^. Rhomboid proteases are part of a broader family of rhomboid proteins, which includes inactive pseudoproteases that perform non-proteolytic functions, such as retrotranslocation of integral membrane proteins for proteasomal degradation ^63^. Given this, we investigated whether the generation of the lumenal fragment of RFP-Vps10-NG-AID involved Rbd2’s proteolytic activity. The predicted catalytic site (Figure 4D) is comprised of a serine (S124) and histidine (H179) dyad, located on TMD4 and TMD6 and is part of a catalytic cleft formed by the last three TMDs, buried within the membrane and open to the luminal side ^61^. We found that GFP-tagged wildtype but not catalytic inactive (S124A) Rbd2, rescued cleavage of the lumenal domain of RFP-Vps10-NG-AID in *rbd2*Δ cells (Figure 4B). Thus, Rbd2 functions as a protease in Vps10 degradation. Imaging of Rbd2-GFP confirmed that it localized to endosomes that partially overlap with TGN and endosomal makers Sec7 and Vps10 (Figure 4E), which is consistent with its previously reported localization ^46, 64^.

Collectively these data demonstrate that Rbd2 is an active rhomboid protease, that localizes to post-ER compartments. It cleaves ubiquitinated RFP-Vps10-NG-AID, releasing soluble lumenal fragments.

### Combined inhibition of the vacuolar proteases, the proteasome, Ddi1 and Rbd2 rescues K48-polybubiquitinated Vps10 from degradation

Our data showed that proteasomes, vacuolar proteases, Ddi1 and Rbd2 each contributed to degradation of K48-polyubiquitinated RFP-Vps10-NG-AID. However, since loss of any single component was insufficient to block degradation, we examined whether degradation could be blocked by their combined loss. Cells with inactivated vacuolar proteases (*pep4*Δ), inactivated proteasome subunits (*pup1-T30A, pre3-T20A*) and sensitized to MG132 (*pdr5*Δ), were knocked out for *DDI1*, *RBD2*, or both.

Combined loss of Rbd2 and Ddi1 along with blocking proteasomes with MG132 prevented formation of any truncated fragments. Instead, a high molecular weight product formed that remained in the SDS-PAGE stacking gel (Figure 5A). The total amount of high MW product was roughly equivalent to the starting amount of full-length protein, although precise measurements were not possible due to the slow migration of the proteins. This high-molecular weight product likely corresponds to highly ubiquitinated RFP-Vps10-NG-AID. In previous experiments, high molecular weight K48 polyubiquitinated proteins also accumulated with loss of Ddi1 ^50, 51^, and were similarly observed to accumulate in the *ddi1*Δ and *ddi1*Δ *rbd2*Δ strains analyzed here (Supplemental Figure 5A).

**Figure 5.**
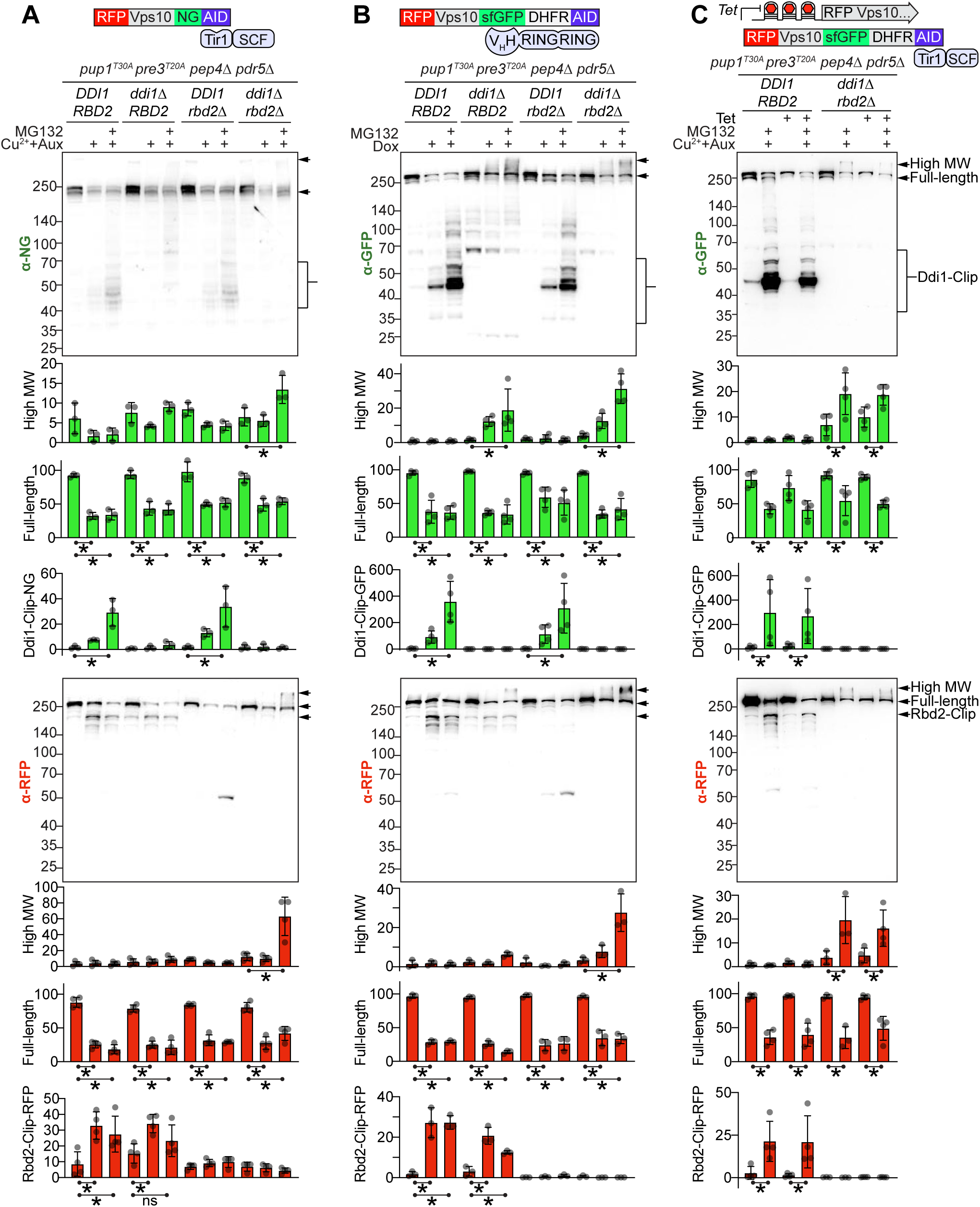
Vps10 is sheared and degraded by Ddi1, Rbd2, the proteasome and vacuolar proteases after K48-linked polyubiquitination **(A-C)** The contribution of Ddi1, Rbd2 and the proteasome to degradation of the indicated ubiquitinated Vps10 fusion proteins was assessed in *pup1-T30A, pre3-T20A pep4*Δ *pdr5*Δ cells lacking either Ddi1, Rbd2, or both. High molecular weight, full-length and clipped forms were monitored by immunoblotting of N-terminal (RFP) or C-terminal tags (NG) and were quantified and graphed (n ≥3, *p<0.05). **A)** RFP-Vps10-NG-AID degradation was induced by SCF^Tir1^. **B)** RFP-Vps10-sfGFP-DHFR-AID degradation was induced by expressing the GFP nanobody RNF4 RING Ub ligase. **C)** RFP-Vps10-sfGFP-DHFR-AID degradation was induced by SCF^Tir1^ 30 min after translation of newly synthesized RFP-Vps10-sfGFP-DHFR-AID was blocked with tetracycline and proteasome activity was blocked with MG132.

These data also indicated that Ddi1, Rbd2 and proteasomes can work independently to target their substrate. Loss of Rbd2 did not impact generation of the Ddi1-dependent cytosolic fragment of RFP-Vps10-NG-AID upon SCF^TIR1^-mediated ubiquitination (Figure 5A). Similarly, loss of Ddi1 did not impact generation of the Rbd2-dependent lumenal fragment. However, when both Ddi1 was inactivated and proteasomes were inhibited, generation of the Rbd2-lumenal product was variable. Rbd2 remained active however, since the full-length protein was degraded and the high molecular weight product did not accumulate. It was surprising that inhibiting proteasomes or deleting *DDI1* did not prevent Rbd2 activity, since if Ddi1 or proteasomes remove the ubiquitinated tail of Vps10, Rbd2 should not be able to recognize the substrate. This could be explained if Rbd2 outcompetes Ddi1 and proteasomes and cleaves the substrate first, alternatively Rbd2 may recognize not only ubiquitinated substrates, but also partially cleaved protein.

The combined actions of Rbd2, Ddi1, and the proteasome were confirmed using an additional K48-ubiquitination system (Figure 5B). Here we used the RNF4-RING Ub ligase and directed it to RFP-Vps10-sfGFP-DHFR-AID, which was cytosolically tagged with superfolder GFP (sfGFP) ^65^ and *E. coli* DHFR ^66^. This tag was used since GFP alone was highly sensitive to proteasomal degradation once targeted by K48-Ub ligases even when proteasome catalytic subunits were inactivated by use of a *pup1-T30A, pre3-T20A* mutant and MG132 (Supplemental Figure 5B-C). Induction of RNF4-RING with doxycycline induced the degradation of full-length RFP-Vps10-sfGFP-DHFR-AID while producing an Rbd2-dependent lumenal fragment and a Ddi1-dependent cytosolic fragment. The combined loss of Rbd2 and Ddi1, along with proteasome inhibition, also resulted in the accumulation of a high molecular weight species at levels comparable to the initial Vps10-sfGFP-DHFR-AID levels before ubiquitination.

Finally, we verified that the degradation products were formed from post-ER Vps10 by preventing the synthesis of new Vps10 during the assay. A tetracycline-sensitive riboswitch ^23, 66^ and a medium strength constitutive promoter was installed upstream of RFP-Vps10-sfGFP-DHFR-AID. This riboswitch binds tetracycline to adopt a secondary structure that prevents translation. RFP-Vps10-sfGFP-DHFR-AID levels slightly decreased after adding tetracycline, suggesting there was some background turnover (Figure 5C). Nonetheless, both Ddi1-dependent and Rbd2-dependent fragments were produced upon SCF^TIR1^-mediated ubiquitination and loss of both Ddi1 and Rbd2 resulted in the accumulation of high molecular weight products upon proteasome inhibition.

Collectively, these data demonstrate that Rbd2, Ddi1, the proteasome and vacuolar proteases independently cleave post-ER Vps10, that has been targeted by K48 ubiquitination.

### Ddi1 and Rbd2 cleave various integral endosomal proteins targeted by K48-Ub ligase

We also examined whether other single pass integral membrane proteins in the endocytic system degraded if they were targeted by K48 Ub-ligases, and if so, whether Ddi1 or Rbd2 targeted these substrates. To monitor for protein shearing, we tagged the N and C terminus of several proteins with RFP and NG-AID, respectively. The proteins occupied different areas of the secretory endomembrane system, but were not always at their reported native localization (Figure 6A). Within 1h of SCF^Tir1^-mediated ubiquitination, all examined proteins degraded, and were partially stabilized by the combined inactivation of Rbd2, Ddi1, the proteasome and vacuolar proteases (Figure 6B). Ylr001c, which was localized to the vacuole limiting membrane, and Ynd1, which was localized to endosomes and the vacuole lumen, both produced a Rbd2-dependent degradation product. They also produced a high molecular weight product upon combined inactivation of Ddi1 and Rbd2. Pex3 and Fet5, which were localized to the ER and vacuole limiting membrane, produced a Ddi1-dependent fragment. Pex3, but not Fet5 produced a high molecular weight product upon Ddi1 inactivation. This indicates that Rbd2 and Ddi1 can cleave multiple ubiquitinated integral membrane substrates throughout the secretory system.

**Figure 6.**
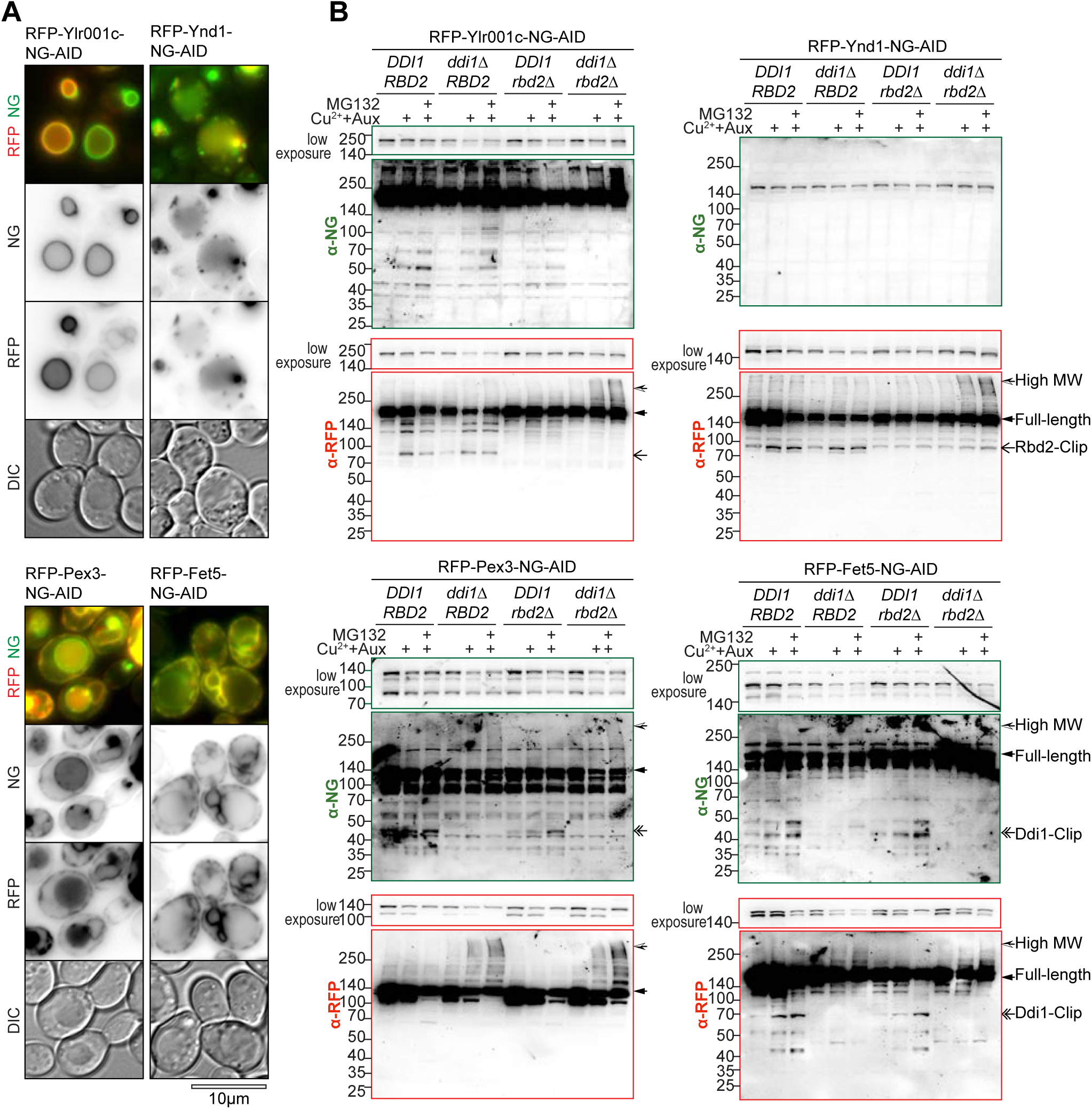
Rbd2 and Ddi1-dependent proteolysis of other membrane proteins in the secretory pathway **A)** Localization and **B)** Immunoblot analysis of lumenal (α-RFP) and cytosolic (α-NG) domains of membrane proteins ubiquitinated by of SCF^Tir1^ in *pup1-T30A, pre3-T20A pep4*Δ *pdr5*Δ cells lacking Rbd2, Ddi1, or both. Bar=10μm.

## Discussion

Our study reveals that the configuration of a Ub modification determines distinct degradative fates for post-ER proteins (Figure 7). Consistent with many previous findings, mono-Ub and K63-linked polyubiquitinated proteins engage MVB sorting and lysosomal degradation ^3, 67^. In contrast K48-polyubiquitinated proteins can engage a protein shearing pathway, we call CUT-UP, which stands for Cleavage of Ubiquitinated Targets by Ubiquitin-activated Proteases (Figure 7). In the CUT-UP pathway, K48-polyubiquitinated proteins are cleaved into smaller fragments for lysosomal and proteasomal degradation. This required proteases that recognize ubiquitinated substrates, which were the proteasome, the intramembrane protease Rbd2, and the cytosolic protease Ddi1.

**Figure 7.**
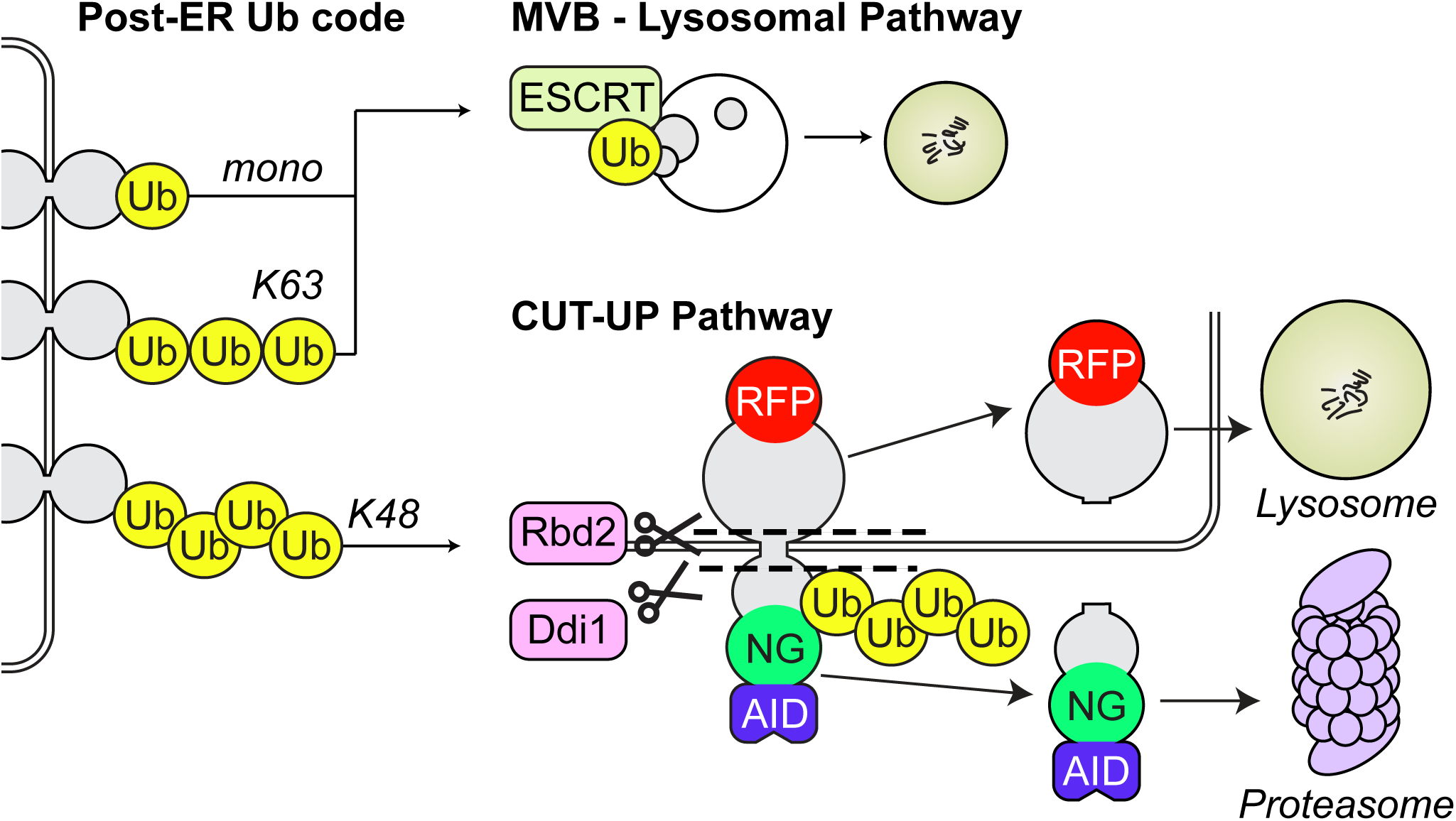
Model for Ub-activated degradation of endocytic cargo Mono-Ub or K63-linked polyubiquitination directs cargo towards MVB sorting and lysosomal degradation. Whereas K48-polyubiquitination directs cargo towards degradation by CUT-UP, a protein shearing pathway. Ubiquitinated proteins are cleaved into smaller fragments for lysosomal and proteasomal degradation by Rbd2, Ddi1, and the proteasome.

Protein shearing might contribute important functions to proteostasis by enabling rapid inactivation of proteins, easier extraction from the membrane (especially for hard to unfold proteins and aggregates), or by serving as a backup degradation pathway when others are compromised ^4, 43, 68, 69^. Protein shearing can also process precursor proteins into active fragments, such as the Ddi1-dependent generation of transcription factors NFR1 and AMOT in human cells, or the Rbd2-dependent generation SREBP in *S. pombe* ^4, 51, 52, 68^. In contrast to other membrane proteases, which recognize peptide sequences that can become exposed with unfolding, Rbd2 and Ddi1 cleave ubiquitinated cargo. This allows their activity to be inducibly activated and deployed within the larger Ub-dependent quality control system.

The CUT-UP pathway required K48-polyubiquitination, which occurs on post-ER membrane proteins and can be mediated by ligases involved in degradation of a wide set of post-ER membrane proteins (Supplementary Discussion) ^2, 70^. This modification might be more prevalent during conditions such as cell stress, whereas ligand-induced downregulation is understood to mostly involve K63-Ub and the MVB pathway.

Interestingly, the EGAD pathway that mediates Cdc48-dependent extraction of proteins from the lipid bilayer was not engaged by K48-polyubiquitination here. This indicates that substrate selection for EGAD involves additional factors, and that perhaps an interaction with the Dsc complex is required before K48-polyubiquitination can facilitate extraction. K48-polyubiquitination is also often used for therapeutic targeted protein degradation. Since shearing might produce toxic fragments, consideration should be given to the degradation mechanism, and to use of an alternative Ub linkage.

Revealing the *in vivo* proteolytic activity of Ddi1 allowed us to explore its mechanism of action. The Ub-dependent proteolytic activity of Ddi1 is contained within its catalytic core, the HDD-RVP domain ^50, 71^, which we found binds mono-Ub directly. We hypothesize that this binding might activate the RVP domain by repositioning flaps that could gate its catalytic site or wrap around the substrate ^45, 55^. The HDD contributes to proteolysis through an unresolved mechanism that involves its C-terminal helix, which, when mutated, inactivates Ddi1. The activity of the HDD-RVP domain is enhanced by auxiliary Ub-binding modules: an atypical UBL and UBA. Since the Ub-binding modules vary across taxa, no specific auxiliary module or linker composition that connects them to the HDD-RVP core seem uniquely essential. Binding assays show Ddi1 prefers K48 over K63-polyUb ^50, 72^ and might be inhibited by some Ub configurations ^71^. However, how Ddi1 achieves K48-polyUb selectivity, particularly *in vivo*, remains unclear, as all its Ub-binding modules, including the HDD-RVP core, bind mono-Ub ^45, 54, 55^.

Deletion of Ddi1 increases the abundance of high-molecular-weight K48-polyubiquitinated proteins, suggesting it has a broad range of substrates ^50, 73^. The only other known function of Ddi1 proteolysis is to repair DNA-protein cross links, but if this involves ubiquitin is unresolved ^74^. The two known substrates of human Ddi2, NRF1 and AMOT, aid in cancer survival ^51, 52, 75–78^. Consequently, there is interest in finding new enzymatic inhibitors to Ddi1. Given that human Ddi1 is relatively resistant to existing HIV protease catalytic site inhibitors, our finding that I183A and D184A mutations in the HDD domain abolish enzymatic activity highlights additional surfaces on Ddi1 that could serve as potential drug targets (Supplemental Figure 3C).

Rbd2 is one of two rhomboids in *S. cerevisiae*. While its paralog, Rbd1, functions as an inner mitochondrial membrane protease that is orthologous to mammalian PARL ^79^, Rbd2’s role was previously unknown. Our study revealed that Rbd2 can cleave a ubiquitinated integral membrane proteins in TGN and vacuolar membranes. A human homolog of Rbd2 is RHBDL4 ^46, 60^. RHBDL4 is one of four rhomboid proteases that occupy the secretory/endosomal system. RHBDL4 contributes to ERAD by cleaving aggregation-prone misfolded proteins, it can also cleave APP and SREBP ^60, 69, 80–82^.

Likewise, *S. pombe* Rbd2 also cleaves SREBP. RHBDL4 is of clinical interest since it is involved in hepatic function, Alzheimer’s progression, and immune dysfunction and is linked to cancer ^82–86^. Both RHBDL4 can bind Cdc48 *via* a SHP box, providing a way to associate with ubiquitinated substrates. RHBDL4 also has a Ub-interaction motif (UIM) that contributes to substrate recognition. These motifs are not present in *Sc* Rbd2 suggesting it uses an alternative intrinsic Ub-binding domain or a Ub-binding adaptor protein.

In conclusion, this study advances three principles of Ub signaling: (1) polyUb linkage determines distinct degradative pathways; (2) secretory membrane proteins undergo Ub-dependent degradation *via* ERAD, MVB and protein shearing pathways; (3) proteases beyond the proteasome can process ubiquitinated substrates. The ubiquitination tools developed here help map the ubiquitin signaling network. Further investigation into the CUT-UP pathway will deepen our understanding of proteostasis.

## Supporting information

Supplemental Discussion

## Acknowledgements

We acknowledge the University of Iowa personnel and instrumentation in the IIHG Genomic Sequencing, the Carver College of Medicine NMR, and Protein & Crystallography core facilities. This work was supported NIH RO1GM058202 to RCP. AYM was supported by ADA postdoctoral fellowship.

Author contributions:AYM, RCP conceptualization; AYM, LY, RCP investigation; AYM, LY visualization; AYM, LY, RCP methodology; AYM writing–original draft; AYM, RCP writing–review and editing; RCP supervision; AYM, RCP funding acquisition.

## Supplemental Tables and Figure Legends

**Supplemental Table 1.** Plasmids used in the study

**Supplemental Table 2.** Yeast Strains used in the study

**Supplemental Table 3.** Table of Antibodies used in the study

**Supplemental Figure 1.**
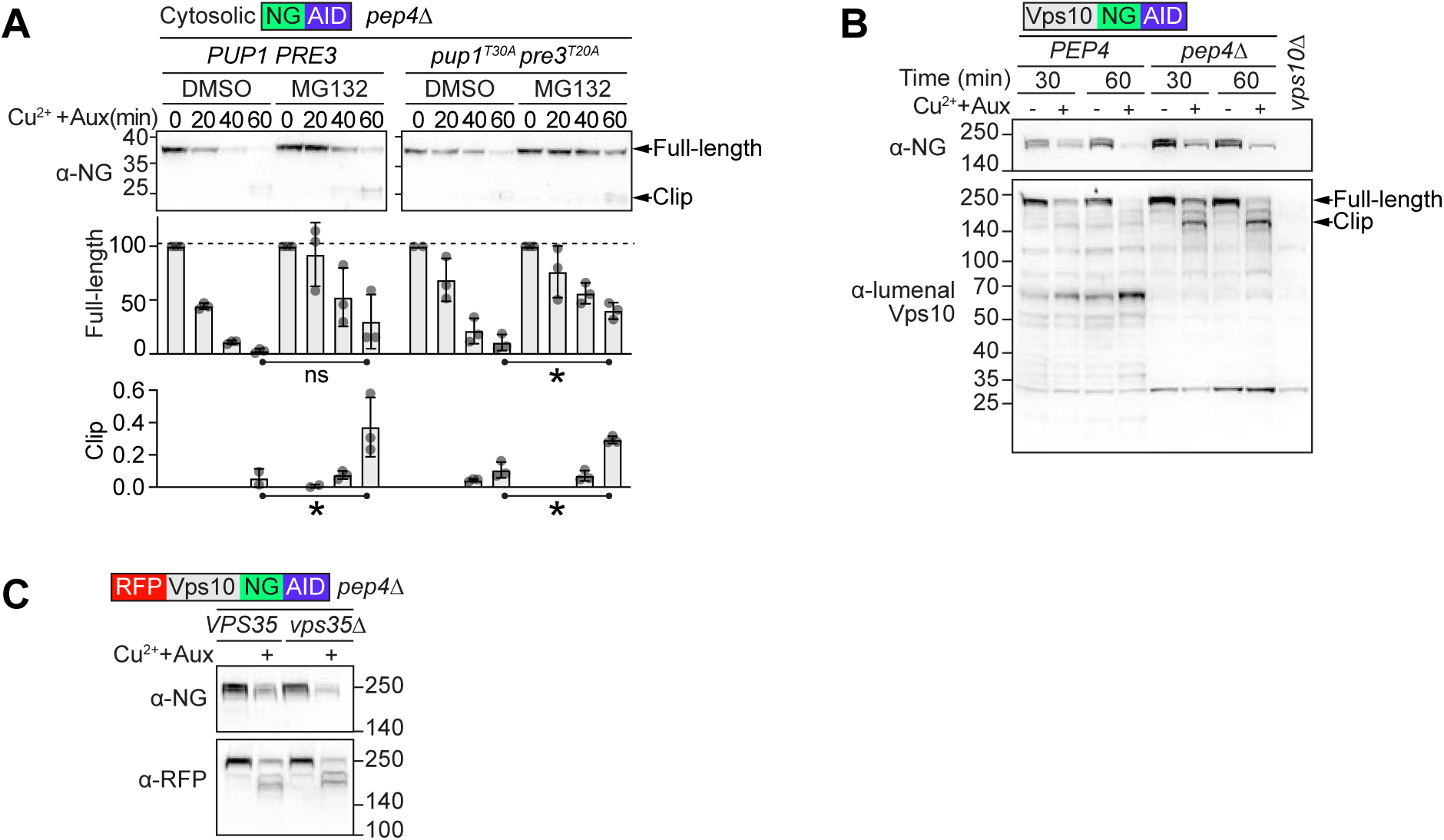
Proteasome inhibition by MG132 and Vps35 and Pep4-independent degradation of Vps10-NG-AID **A)** SCF^Tir1^ dependent degradation of a cytosolic protein (NG-AID) was monitored in *PUP1 PRE3* or *pup1-T30A pre3-T20A* cells. MG132 was added 30 min prior to induction of SCF^Tir1^ with copper and auxin. Representative immunoblot and graph of full-length and clipped fragments are shown (n ≥3, *p<0.05). **B)** Immunoblot of SCF^Tir1^ dependent degradation of Vps10-NG-AID in *PEP4* and *pep4*Δ cells using α-NG and α-Vps10 antibodies. **C)** Immunoblot of RFP-Vps10-NG-AID degradation in *VPS35* and *vps35*Δ cells.

**Supplemental Figure 2.**
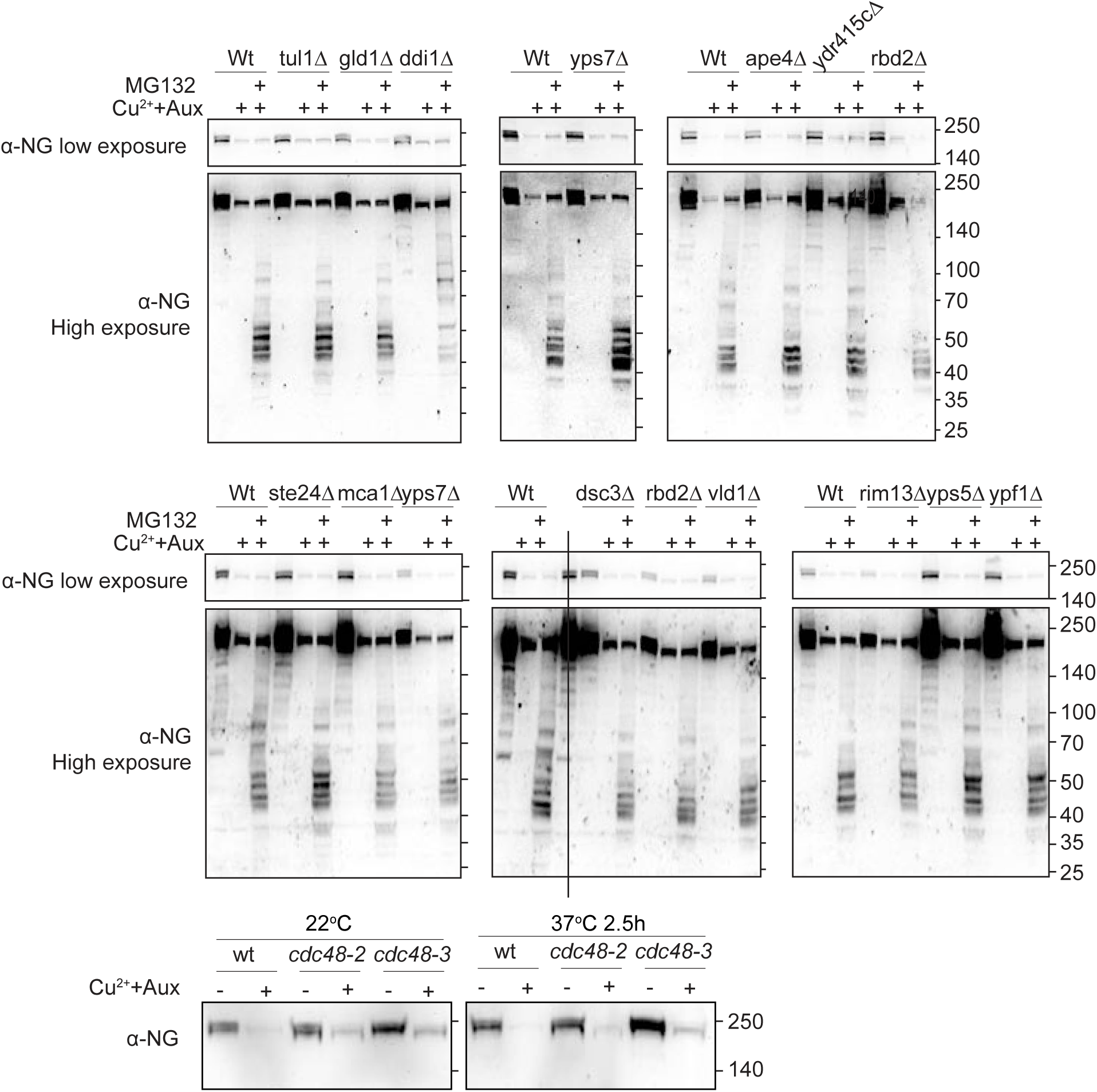
Genetic screen for mediators of Vps10-NG-AID degradation Immunoblot of SCF^Tir1^-dependent degradation of Vps10-NG-AID degradation in the indicated genetic mutants. MG132 was added 30 min prior to induction of SCF^Tir1^ with copper and auxin. A low and high exposure of the same immunoblot using α-NG antibody is shown. Cdc48 temperature-sensitve mutants were grown at 22°C and then shifted to 37°C mutants for 2.5 h at 37°C prior to induction of SCF^Tir1^-dependent degradation of Vps10-NG-AID.

**Supplemental Figure 3.**
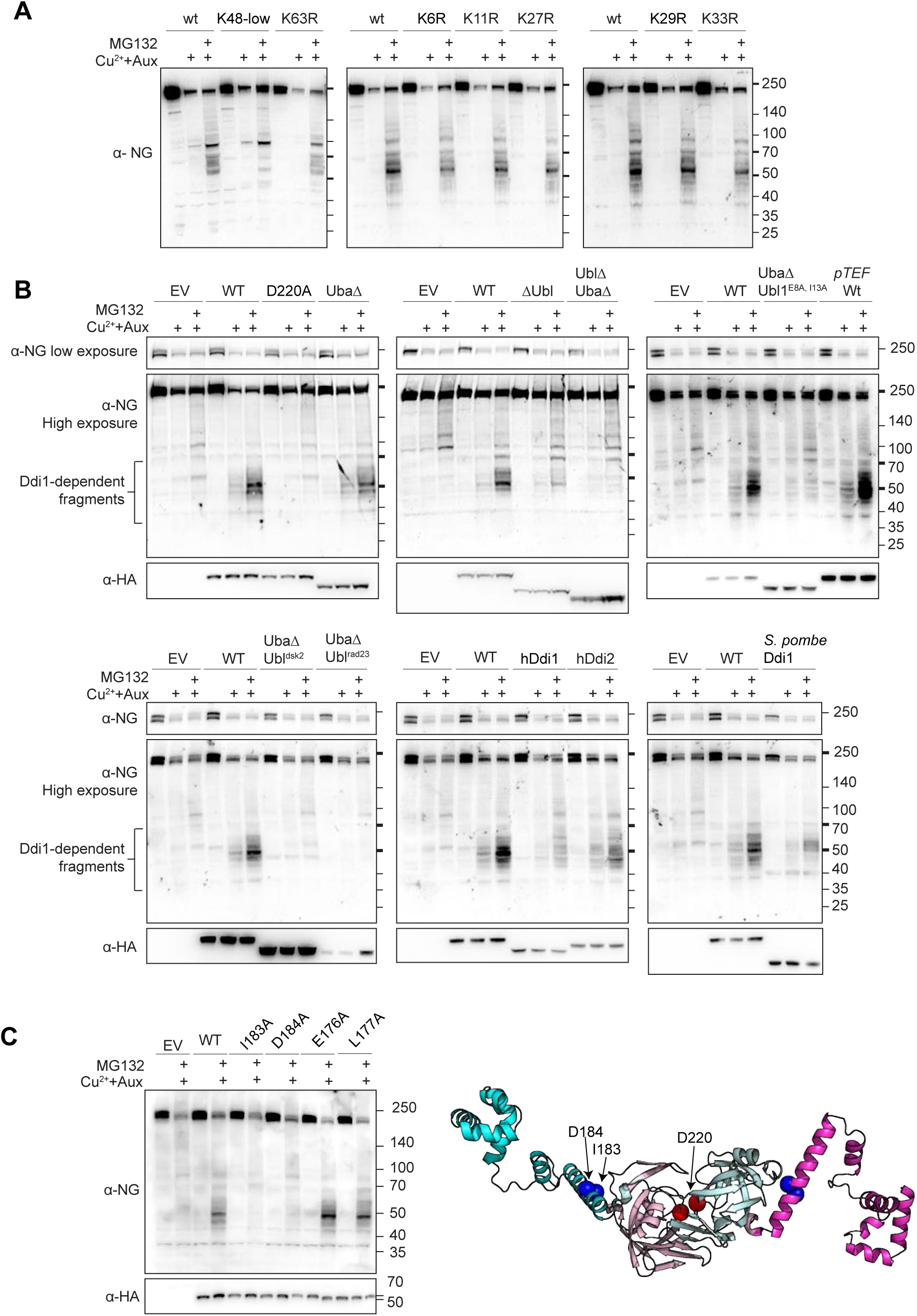
Ddi1-dependent degradation of Vps10-NG-AID **A)** Representative immunoblot of Vps10-NG-AID degradation in strains carrying the indicated K>R Ub mutants, except for K48R Ub, which also carries low levels of wildtype Ub to sustain viability. These data were quantified and presented in Figure 3B. **B)** Representative immunoblots of Vps10-NG-AID degradation in *pup1-T30A pre3-T20A ddi1*Δ cells expressing Ddi variants that were quantified and presented in Figure 3C. **C)** *Left,* immunoblot of Vps10-NG-AID *pup1-T30A pre3-T20A ddi1*Δ cells expressing Ddi variants with mutations in HDD domain. *Right,* Alphafold schematic of HDD-RVP domain of Ddi1 as a dimer, with inactivating mutations in HDD highlighted blue and catalytic site highlighted red.

**Supplemental Figure 4.**
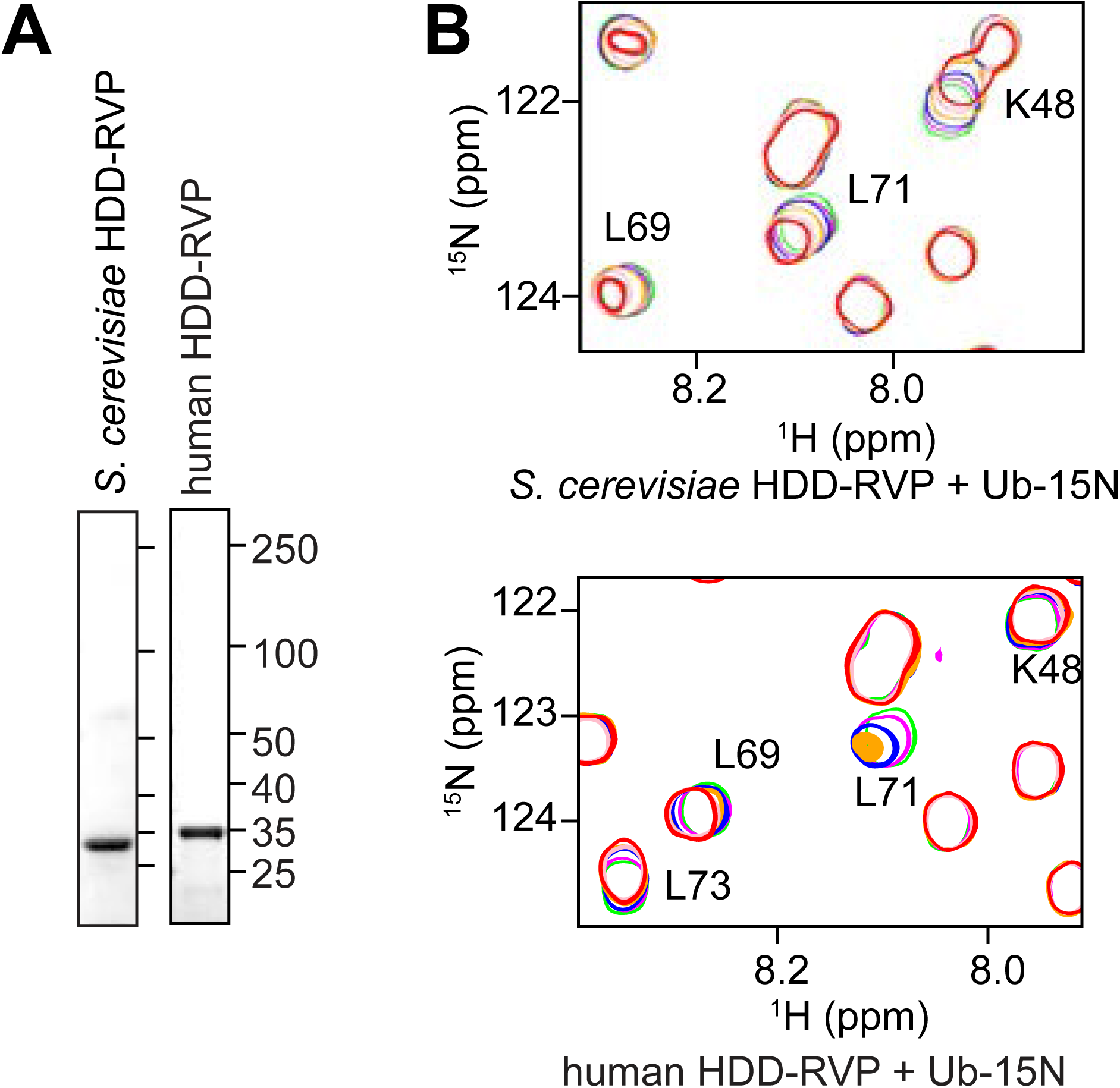
HSQC analysis of ^15^N-Ub in the presence of the HDD-RVP domains from *Sc*Ddi1 and human DDI2 **A)** SDS-PAGE and Coomassie staining of purified HDD-RVP domains. **B)** Portion of the overlay of the HSQC spectra of 30µM ^15^N-Ub at various concentrations of HDD-RVP domain of *Sc*Ddi1 and human DDI2 following the color code in Figure 3D-E.

**Supplemental Figure 5.**
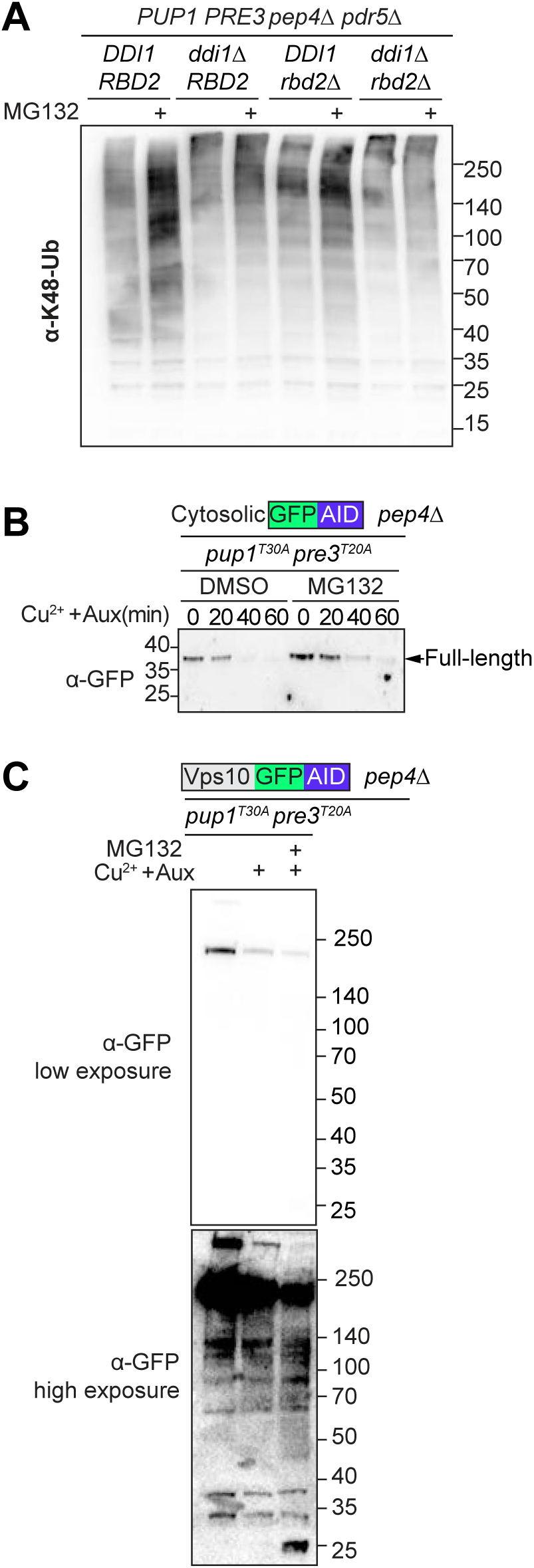
K48-Ubiquitinated proteins in Ddi1 mutants and SCF^Tir1^ mediated degradation of GFP **A)** Immunoblot of whole cell lysates of K48-Ub in *PUP1 PRE3 pep4*Δ *pdr5*Δ cells lacking Ddi1, Rbd2 or both using α-K48-polyUb antibody. MG132 was added for 1 h prior to lysis. **(B-C)** Immunoblot of (α-GFP) of GFP-AID **(B)** or Vps10-GFP-AID **(C)** expressed from the *Vps10* promoter in *pup1-T30A, pre3-T20A pep4*Δ *pdr5*Δ cells. MG132 was added 30 min prior to induction of SCF^Tir1^ with copper and auxin. GFP-AID was only slightly stabilized by MG132 in contrast to NG-AID observed in Supplemental Figure 1A. Stabilization of a C-terminal fragment of Vps10-GFP-AID was also observed upon proteasome inhibition, but had low abundance likely due to vulnerability of GFP to proteasome degradation.

Supplemental Discussion. References documenting K48-ubiquitination or proteasomal degradation of post-ER proteins.

