## Supplemental Discussion for "K48-ubiquitin-activated proteases cut-up post-ER proteins"

**Supplemental Discussion.** References documenting K48-ubiquitination or proteasomal degradation of post-ER proteins.

1. Lin H, Gao D, Hu MM, Zhang M, Wu XX, Feng L, Xu WH, Yang Q, Zhong X, Wei J, Xu ZS, Zhang HX, Song ZM, Zhou Q, Ye W, Liu Y, Li S, Shu HB. **MARCH3 attenuates IL-1beta-triggered inflammation by mediating K48-linked polyubiquitination and degradation of IL-1RI.** Proc Natl Acad Sci U S A. 2018;115(49):12483-8. Epub 2018/11/18. doi: 10.1073/pnas.1806217115. PubMed PMID: 30442668; PMCID: PMC6298087.

- K48 ubiquitination of IL-1 receptor 1 by MARCH3.

2. Chen R, Li M, Zhang Y, Zhou Q, Shu HB. **The E3 ubiquitin ligase MARCH8 negatively regulates IL-1beta-induced NF-kappaB activation by targeting the IL1RAP coreceptor for ubiquitination and degradation.** Proc Natl Acad Sci U S A. 2012;109(35):14128-33. Epub 2012/08/21. doi: 10.1073/pnas.1205246109. PubMed PMID: 22904187; PMCID: PMC3435212.

- Overexpression of MARCH8, which localizes to endoscomes and the plasma membrane, led to K48 polyubiquitination and degradation of IL1RAP.

3. Lu JC, Piazza TM, Schuler LA. **Proteasomes mediate prolactin-induced receptor down-regulation and fragment generation in breast cancer cells.** J Biol Chem. 2005 Oct 7;280(40):33909-16. doi: 10.1074/jbc.M508118200. Epub 2005 Aug 15. PMID: 16103113; PMCID: PMC1976473.

- Ub-dependent shearing of the Prolactin Receptor.

4. Varghese B, Barriere H, Carbone CJ, Banerjee A, Swaminathan G, Plotnikov A, Xu P, Peng J, Goffin V, Lukacs GL, Fuchs SY. **Polyubiquitination of prolactin receptor stimulates its internalization, postinternalization sorting, and degradation via the lysosomal pathway.** Mol Cell Biol. 2008;28(17):5275-87. Epub 2008/06/25. doi: 10.1128/MCB.00350-08. PubMed PMID: 18573876; PMCID: PMC2519723.

- Ligand stimulated both K48- and K63 polyubiquitination of prolactin receptor, the latter was involved in endocytosis.

5. Kumar KG, Barriere H, Carbone CJ, Liu J, Swaminathan G, Xu P, Li Y, Baker DP, Peng J, Lukacs GL, Fuchs SY. **Site-specific ubiquitination exposes a linear motif to promote interferon-alpha receptor endocytosis.** J Cell Biol. 2007 Dec 3;179(5):935-50. doi: 10.1083/jcb.200706034. PMID: 18056411; PMCID: PMC2099190.

- SCFβTrCP ubiquitinates interferon (IFN)-α/β receptor 1 (IFNAR1) subunit of the type I IFN receptor with K63- and K48-polyubiquitination.  K48-ubiquitination is required for maximal degradation of IFNAR1.

6. Feng L, Li C, Zeng LW, Gao D, Sun YH, Zhong L, Lin H, Shu HB, Li S. **MARCH3 negatively regulates IL-3-triggered inflammatory response by mediating K48-linked polyubiquitination and degradation of IL-3Rα.** Signal Transduct Target Ther. 2022 Jan 24;7(1):21. doi: 10.1038/s41392-021-00834-7. PMID: 35075102; PMCID: PMC8786845.

- Cell surface localizes MARCH3 mediates ligand-stimulated K48-polyubiquitination of  IL-3Rα.

7. Zhu LL, Luo TM, Xu X, Guo YH, Zhao XQ, Wang TT, Tang B, Jiang YY, Xu JF, Lin X, Jia XM**. E3 ubiquitin ligase Cbl-b negatively regulates C-type lectin receptor-mediated antifungal innate immunity.** J Exp Med. 2016;213(8):1555-70. Epub 2016/07/20. doi: 10.1084/jem.20151932. PubMed PMID: 27432944; PMCID: PMC4986534.

- Cbl-b facilitates the K48-polyubiquitination of Dectin-2 or Dectin-3 after activation by α-mannans on the surfaces of C. albicans hyphae.

8. Sehat B, Andersson S, Girnita L, Larsson O. **Identification of c-Cbl as a new ligase for insulin-like growth factor-I receptor with distinct roles from Mdm2 in receptor ubiquitination and endocytosis.** Cancer Res. 2008;68(14):5669-77. Epub 2008/07/18. doi: 10.1158/0008-5472.CAN-07-6364. PubMed PMID: 18632619.

- Upon ligand stimulation, c-Cbl associates with cell surface IGF-IR and mediates receptor K48-polyubiquitination.

9. Li Y, Kumar KG, Tang W, Spiegelman VS, Fuchs SY. **Negative regulation of prolactin receptor stability and signaling mediated by SCF(beta-TrCP) E3 ubiquitin ligase.** Mol Cell Biol. 2004;24(9):4038-48. Epub 2004/04/15. doi: 10.1128/MCB.24.9.4038-4048.2004. PubMed PMID: 15082796; PMCID: PMC387770.

- Upon ligand stimulation, SCFβTrCP associates with PRLR, ubiquitinates it and enhances its degradation.

10. van Kerkhof P, Westgeest M, Hassink G, Strous GJ. **SCF(TrCP) acts in endosomal sorting of the GH receptor. Exp Cell Res.** 2011;317(7):1071-82. Epub 2011/01/05. doi: 10.1016/j.yexcr.2010.12.020. PubMed PMID: 21195069.

- GHR is targeted by SCFβTrCP and targeted to MVB pathway.

11. van Kerkhof P, Alves dos Santos CM, Sachse M, Klumperman J, Bu G, Strous GJ. **Proteasome inhibitors block a late step in lysosomal transport of selected membrane but not soluble proteins**. Mol Biol Cell. 2001 Aug;12(8):2556-66. doi: 10.1091/mbc.12.8.2556. PMID: 11514635; PMCID: PMC58613.

- Proteasome inhibitors block degradation of truncated GHR lacking lysine residues

12. Meijer IM, van Rotterdam W, van Zoelen EJ, van Leeuwen JE. **Cbl and Itch binding sites in ERBB4 CYT-1 and CYT-2 mediate K48- and K63-polyubiquitination, respectively**. Cell Signal. 2013;25(2):470-8. Epub 2012/11/17. doi: 10.1016/j.cellsig.2012.11.008. PubMed PMID: 23153581.

- Chimera of EGFR-ERBB4 receptor is K48 and K63-polyubiqutinated.

13. Li Y, Zhou Z, Alimandi M, Chen C. **WW domain containing E3 ubiquitin protein ligase 1 targets the full-length ErbB4 for ubiquitin-mediated degradation in breast cancer**. Oncogene. 2009;28(33):2948-58. Epub 2009/06/30. doi: 10.1038/onc.2009.162. PubMed PMID: 19561640.
    - ERBB4-Cyt1 isoform is partly degraded by proteasomes after ubiquitinated by WWP1.

14. Hoppe T, Matuschewski K, Rape M, Schlenker S, Ulrich HD, Jentsch S. **Activation of a membrane-bound transcription factor by regulated ubiquitin/proteasome-dependent processing.** Cell. 2000 Sep 1;102(5):577-86. doi: 10.1016/s0092-8674(00)00080-5. PMID: 11007476.

- Spt23 and Mga2 are single-pass integral membrane proteins in the ER, which are cleaved into transcription factors by the proteasome.

15. Walrafen P, Verdier F, Kadri Z, Chrétien S, Lacombe C, Mayeux P. **Both proteasomes and lysosomes degrade the activated erythropoietin receptor**. Blood. 2005 Jan 15;105(2):600-8. doi: 10.1182/blood-2004-03-1216. Epub 2004 Sep 9. PMID: 15358619.

- EpoR is jointly degraded by the proteasome and lysosome

16. Meyer L, Deau B, Forejtníková H, Duménil D, Margottin-Goguet F, Lacombe C, Mayeux P, Verdier F. **beta-Trcp mediates ubiquitination and degradation of the erythropoietin receptor and controls cell proliferation**. Blood. 2007 Jun 15;109(12):5215-22. doi: 10.1182/blood-2006-10-055350. Epub 2007 Feb 27. PMID: 17327410.

- EpoR is ubiquitinated by K48-Ub ligase SCF-βTrCP and degraded by both the proteasome and lysosome.

17. Kim H, Vick P, Hedtke J, Ploper D, De Robertis EM. **Wnt Signaling Translocates Lys48-Linked Polyubiquitinated Proteins to the Lysosomal Pathway**. Cell Rep. 2015 May 26;11(8):1151-9. doi: 10.1016/j.celrep.2015.04.048. PMID: 26004177.

- K48-polyUb accumulates on endolyosomes after activation of Wnt signalling.

18. Manohar S, Jacob S, Wang J, Wiechecki KA, Koh HWL, Simões V, Choi H, Vogel C, Silva GM. **Polyubiquitin Chains Linked by Lysine Residue 48 (K48) Selectively Target Oxidized Proteins In Vivo.** Antioxid Redox Signal. 2019 Nov 20;31(15):1133-1149. doi: 10.1089/ars.2019.7826. PMID: 31482721; PMCID: PMC6798811.

- Acute treatment of cells with H2O2 leads to possible K48-polyubiquitination of several post-ER membrane proteins in yeast. Ubiquitinated proteins were purified in a K63R strain.

19. Okiyoneda T, Veit G, Sakai R, Aki M, Fujihara T, Higashi M, Susuki-Miyata S, Miyata M, Fukuda N, Yoshida A, Xu H, Apaja PM, Lukacs GL. **Chaperone-Independent Peripheral Quality Control of CFTR by RFFL E3 Ligase**. Dev Cell. 2018 Mar 26;44(6):694-708.e7. doi: 10.1016/j.devcel.2018.02.001. Epub 2018 Mar 1. PMID: 29503157; PMCID: PMC6447300.

- Post-ER ΔF508-CFTR is K48 and K63-ubiquitinated. Proteasome inhibition increases K48-ubiuqitination of post-ER ΔF508-CFTR
